# A pipeline for multidimensional confocal analysis of mitochondrial morphology, function and dynamics in pancreatic β-cells

**DOI:** 10.1101/687749

**Authors:** Ahsen Chaudhry, Rocky Shi, Dan S. Luciani

## Abstract

Live-cell imaging of mitochondrial function and dynamics can provide vital insights into both physiology and pathophysiology, including of metabolic diseases like type 2 diabetes. However, without super-resolution microscopy and commercial analysis software it is challenging to accurately extract features from dense multi-layered mitochondrial networks, such as those in insulin-secreting pancreatic β-cells. Motivated by this, we developed a comprehensive pipeline, and associated ImageJ plugin, that enables 2D/3D quantification of mitochondrial network morphology and dynamics in mouse β-cells, and by extension other similarly challenging cell-types. The approach is based on standard confocal microscopy and shareware, making it widely accessible. The pipeline was validated using mitochondrial photo-labelling and unsupervised cluster analysis, and is capable of morphological and functional analyses on a per-organelle basis, including in 4D (*xyzt*). Overall, this tool offers a powerful framework for multiplexed analysis of mitochondrial state/function, and provides a valuable resource to accelerate mitochondrial research in health and disease.

## INTRODUCTION

Mitochondria are the main energy producing organelle of eukaryotic cells and are essential for a diverse range of cellular functions, including ATP synthesis, Ca^2+^ homeostasis, ROS signaling, and the control of apoptotic cell death (1, 2). Microscopy has been instrumental in unraveling intricacies of mitochondrial biology and their diverse roles in cellular physiology and pathophysiology. Electron microscopy has provided fundamental insights into mitochondrial ultrastructure and cellular distribution in health and disease but requires cell fixation and only provides a static snapshot. In contrast, fluorescence microscopy of live cells, labeled with mitochondria-targeted fluorescent proteins or dyes, has revealed that mitochondria are highly dynamic and motile organelles that undergo frequent fusion and fission events (3–5). Mitochondrial dynamics and network morphology vary in different cellular states, and are important for the function and quality control of the organelle, as well as overall cell health and adaptation to stress (1). Healthy mitochondria are generally mobile, tubular in shape and exist in complex networks, whereas cells undergoing profound stress or entering apoptosis often display swollen and fragmented mitochondria, marked by concurrent disruption of metabolism, membrane potential, ROS levels, and Ca^2+^ signalling (6–8). Quantitative imaging-based assessment of mitochondrial morphology and dynamics can therefore provide valuable insights into cellular physiology and pathophysiology.

In pancreatic β-cells, mitochondria play an essential role in insulin secretion, which relies on ATP and other mitochondria-derived metabolites to both trigger and amplify insulin granule exocytosis in response to glucose and other nutrient stimuli (9). Dysfunction of β-cell mitochondria therefore results in loss of glucose-stimulated insulin secretion (10). Perturbations to mitochondria are also a common feature in insulin target tissues with impaired insulin signaling (11, 12). Mitochondria thus take center stage in both β-cell failure and insulin resistance and are an area of significant focus in efforts to understand the pathophysiology of type 2 diabetes (13–15). Mitochondria also exist as dynamic networks in β-cells. Fusion within the network may help protect β-cells from nutrient stress-induced apoptosis (6), and mitochondrial fragmentation, swelling and dysfunction are seen in β-cells from patients with type 2 diabetes and rodent models of diabetes (15–18). Normal insulin secretion may also be influenced by β-cell mitochondrial dynamics (19–21), but exactly how networking of the organelle relates to its metabolic capacity in healthy β-cells or during conditions of moderate nutrient excess remains unclear and warrants further investigation.

Most types of microscopy can detect the prominent morphological differences between healthy and severely stressed mitochondria with relative ease. It is, however, much more challenging to accurately quantify subtle changes in mitochondrial dynamics, or perform 3D analysis of the full mitochondrial network. This is particularly difficult in cells with a dense mitochondrial network that spans several layers, such as β-cells (6, 18). Although methods have been published that integrate 3D confocal imaging and analysis of mitochondria, these generally use commercial software packages and/or are optimized for relatively flat cell types (22, 23). This is likely one reason why there are only few quantitative analyses of β-cell mitochondrial dynamics, and why full 3D investigations of β-cell mitochondria are limited to a small number of examples using super-resolution approaches, such as 4π-microscopy (18, 24).

To facilitate progress in the important area of mitochondrial biology and dynamics we present here a pipeline for quantitative multidimensional analysis of mitochondria that is based on standard confocal fluorescence microscopy and the open source image analysis platform ImageJ/Fiji (25, 26). In this, we identify a superior method for accurate identification of individual mitochondria within dense networks, and we outline a framework for quantitative description of mitochondrial morphology and network characteristics. Applying this pipeline to clonal MIN6 β-cells and primary mouse β-cells, we quantitatively distinguish mitochondrial morphologies, including the functional and morphological changes to physiological and pathophysiological stimuli. Additionally, we discuss the pros and cons of 2D and 3D imaging approaches, identify image processing steps required for accurate mitochondrial analysis in 3D, and apply these to quantitate distinct 3D β-cell network morphologies. Finally, we extend our analysis to 4D by including time-lapse data, and we demonstrate the feasibility of using the pipeline to quantitate the dynamics of the entire three-dimensional mitochondrial network in live cells.

## RESULTS AND DISCUSSION

### Overall workflow & general considerations

Fluorescence confocal analysis of mitochondria in live cells involves several general steps, each of which is important for high-quality results (Figure 1). As a starting point, the cells must be cultured on glass coverslips, or other vessels, that are appropriate for confocal microscopy. The mitochondria should then be labelled using carefully chosen mitochondria-targeted fluorescent proteins or organic dyes (27), and the image acquisition should be optimized and carried out in a manner that provides sufficiently high resolution and image quality for accurate analysis. Because these factors and general steps can vary between specific experiments and microscope systems, an extensive discussion falls beyond the scope of this paper. The imaging parameters and conditions we have used are detailed in Materials and Methods. Our focus in the following will be on the post-acquisition steps that are critical for accurate morphological analysis of mitochondria in the confocal images.

**Figure 1:**
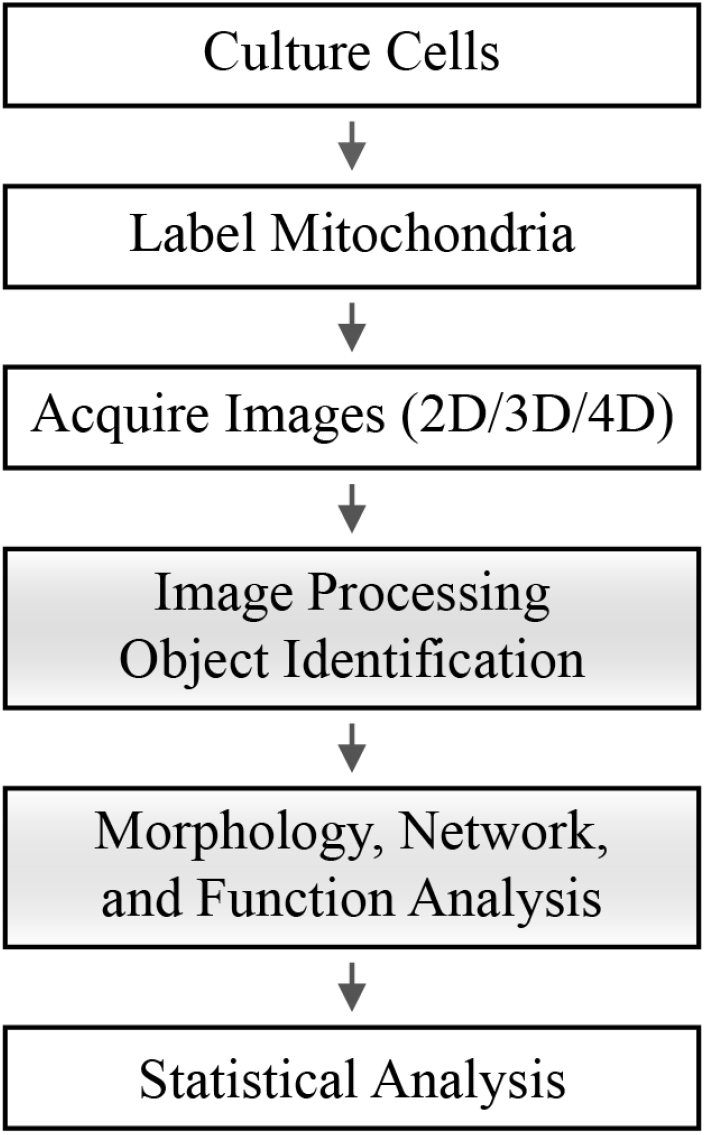
Schematic of the general workflow required for mitochondrial analysis by confocal microscopy. The shaded boxes represent the steps that are addressed and detailed in this paper.

Image acquisition and analysis can be done in 2D or 3D, and by further extending this to include time-lapse capture, important information can be extracted about mitochondrial dynamics. The choice between these imaging modes may be influenced by several considerations, including the type and thickness of the cell, the specific parameters to be quantified, and the biological questions being asked. For instance, we will discuss later how some 2D analyses of relatively thick cells, such as pancreatic β-cells, can be associated with inaccuracies that may be mitigated by a full 3D analysis of the mitochondrial network. In all cases, accurate quantification of mitochondrial features involves image processing steps and identification of the mitochondrial objects in the image. Morphological features can then be extracted using appropriate 2D or 3D shape descriptors, while mitochondrial networking can be assessed through skeletonization analysis. In this latter process, the binarized mitochondria are converted into topological skeletons (the thinnest form that is equidistant to its edges) and the branches of the skeleton are analyzed. In the following, we describe each of these post-acquisition steps and identify a number of “best approaches”, to build a pipeline for accurate multidimensional analysis of mitochondria that we also implement and make available in a comprehensive Mitochondria Analyzer plugin for ImageJ/Fiji (28).

### Image thresholding and identification of mitochondria

Before accurate morphological analysis of fluorescently–labeled mitochondria can be done, it is essential that: i) the mitochondrial population is correctly identified in the images, and ii) the individual mitochondrial units can be distinguished within the dense mitochondrial network. For this critical step, a thresholding process based on analysis of the intensity histogram is used to distinguish pixels with “true” fluorescent signal from background signal. This process also groups any identified positive pixels that are connected into discrete objects (i.e. mitochondria) that can be analyzed further. Thresholding approaches can be broadly categorized as either ‘Global’ or ‘Local’, which identify positive pixels based on the histogram of the entire image or on dynamic analyses of image sub-regions, respectively (22, 29). Global thresholding tends to be the most commonly used approach, but this may reflect its relative ease of use rather than accuracy.

To identify the most suitable thresholding strategy for mitochondrial identification, we compared the performance of the Global and Local threshold methods available for ImageJ/Fiji on images of primary islet cells stained with MitoTracker dye. This was judged on the ability to preserve mitochondrial structural detail while minimizing capture of background signal. For optimal results, all images were pre- and post-processed to reduce noise (see Materials & Methods). Among the Global-based algorithms in the ImageJ/Fiji “Auto Threshold” command, we qualitatively estimated that the Default method performs similar to, or in several cases better than, the other Global algorithms (Figure S1).

The Local thresholding methods we tested included the Mean, Median, and Mid-grey algorithms (part of the “Auto Local Threshold” command), as well as the Weighted Mean method (also called Adaptive threshold), which is available through a separate plugin (30) (Figure S2). These Local methods compute a threshold for each pixel in the image and require the definition of two parameters: a block size and a C-value. The block size specifies the size of the region around each pixel for which the histogram is analyzed and should be chosen based on the size of the objects of interest for the best results (30). The C-value provides an offset to the threshold and helps strike a balance between minimizing noise detection and incorrectly splitting objects into smaller pieces (30, 31). Using the Adaptive threshold method for optimization, we found that the ideal C-value depended on the image’s signal-to-noise contrast and needed to be empirically determined. For each set of images that has been acquired and processed in a similar manner, we therefore recommend that various combinations of block size and C-values should be tested on a representative image to determine the best combination. The optimized parameters can then be used to threshold all images in the group similarly (Figure S2, Supplemental Methods for details). Among the Local threshold approaches, our assessment was that the Mean and Adaptive threshold methods best captured mitochondrial structure, and that the Adaptive threshold further tended to identify less noise (Figure S2B).

A side-by-side comparison indicated that Local (Adaptive) thresholding resolves mitochondria better than Global thresholding, which appears to capture more out-of-focus signal and/or noise, and therefore erroneously merges adjacent objects (Figures 2A & B). For a more stringent and quantitative evaluation, we used mitochondria-targeted photoactivatable GFP (mito-PAGFP) to selectively photo-label single mitochondria and identify truly contiguous organelles within dense regions of the network (4, 5). As exemplified in Figure 2B, and quantified in Figure 2C (see also Figure S3), Adaptive thresholding was indeed better at delineating photo-labeled mitochondria, and distinguishing closely adjacent parts of the network that are physically separate. In contrast, the Global threshold algorithm consistently overestimated the mitochondrial size. Taken together, these comparisons established that using Adaptive thresholding, with empirically optimized parameter values, is a superior approach for accurate identification of fluorescently labeled β-cell mitochondria.

**Figure 2:**
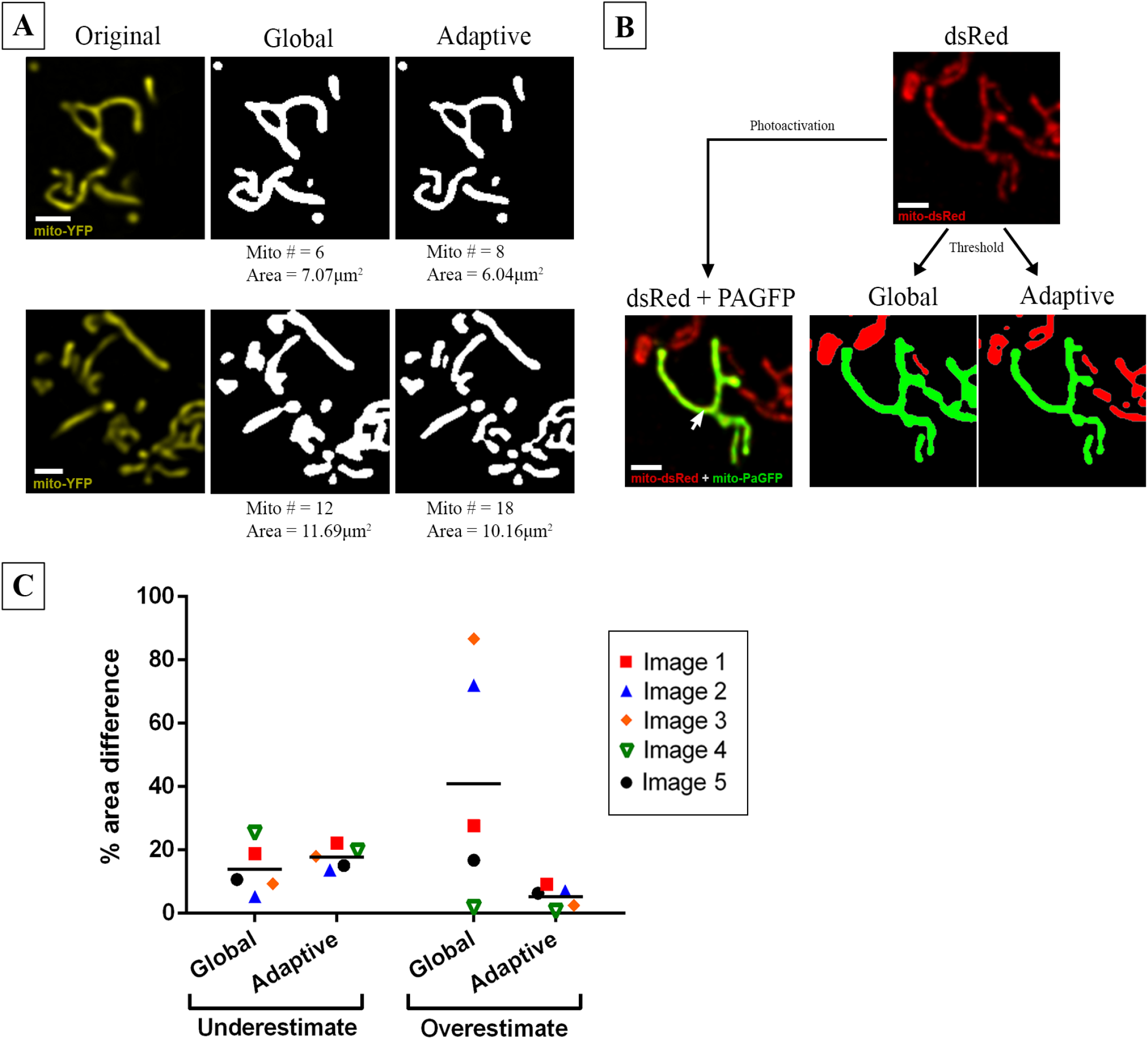
Comparison of mitochondrial identification using Global vs Adaptive thresholding methods. **(A)** Two representative examples of object identification using Global thresholding (‘Default’ method) vs Adaptive thresholding (radius = 1.25 μm, C = 11) on images of MIN6-cell mitochondria labelled with Mito-YFP. The number of identified objects (mitochondria) and their total area are indicated below the images. Scale bar = 1 μm. **(B)** Part of the mitochondrial network in a MIN6 cell co-transfected with mito-dsRed and mito-PAGFP. *Top -* All mitochondria imaged in the mito-dsRed channel. *Bottom left -* A single mitochondrion (green) was labeled by laser-based mito-PAGFP activation at the point indicated by the arrow. *Bottom right -* Object identification using Global vs Adaptive threshold algorithms applied to the dsRed channel; in each image, the object that is identified as contiguous with the PAGFP-labelled mitochondrion is shown in green. Comparison with the original image shows that the Adaptive method more accurately distinguished the photo-labeled mitochondrion, whereas Global thresholding artificially merged it with adjacent mitochondria. Scale bar = 1 μm. (**C**) Quantitative comparison of the degree to which Global and Adaptive thresholding under- or over-estimated the PAGFP-labeled mitochondrion in 5 test images. The corresponding images and details of the estimation algorithm are shown in Figure S3.

### Two-dimensional analysis of mitochondrial morphology and network connectivity

After careful image thresholding, the next step is to quantify the morphological features of the identified mitochondrial objects. We therefore identified a comprehensive set of parameters to capture and mathematically describe key aspects of the mitochondrial morphology. For 2D analysis, we characterize mitochondrial size by area and perimeter, while mitochondrial shape is defined by form factor (FF) and aspect ratio (AR). We evaluate the overall connectivity and morphological complexity of the mitochondrial network based on the skeletonized network, and quantify this by the number of branches, the number of branch junctions, as well as total (accumulated) length of branches in the skeleton. Figure 3 summarizes the various parameters and indicate how they change with various morphologies.

**Figure 3:**
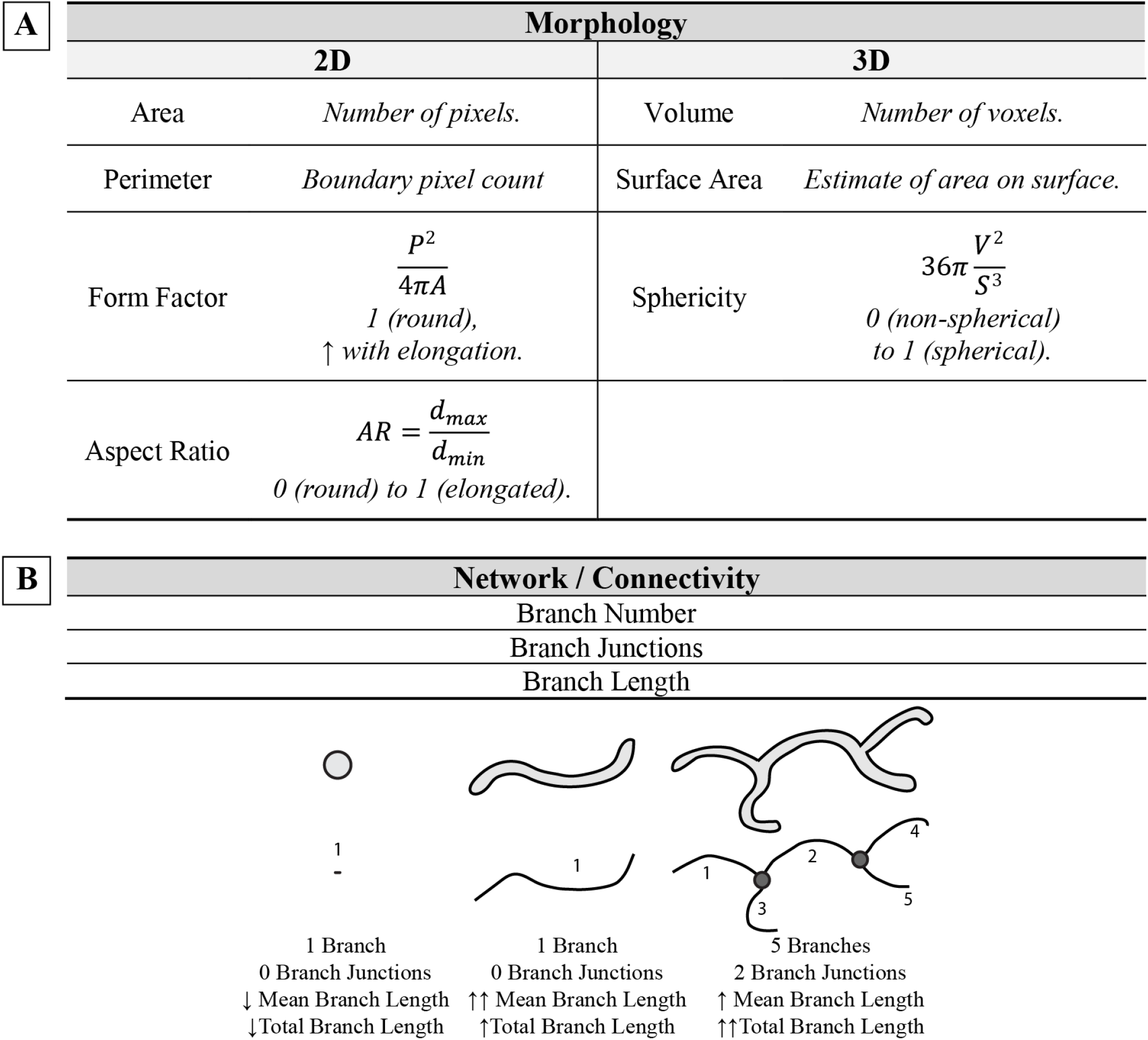
Summary of morphological and network descriptors used in 2D/3D analysis of mitochondria. **(A)** Summary and definitions of the mathematical descriptors used to quantify mitochondrial morphology in two and three dimensions. **(B)** Summary of parameters used to describe mitochondrial network connectivity and illustration of skeletonization analysis on: a punctate object with no branch junctions and minimal branch length (*Left*); a long single tubular object with no branch junctions but higher branch length (*Middle*); and a complex object with multiple branches and junctions (gray dots), and the highest total branch length (*Right*). Note that the mean branch length, derived by dividing the total branch length by number of branches, is greater in the object in the middle than the more complex object on the right.

To evaluate the ability of this approach to measure and distinguish mitochondrial morphologies, we transfected MIN6 cells with mito-YFP and generated an image-set consisting of 2D slices from 84 cells. We then divided the cells into three different categories based on visual inspection of their mitochondria: 1) a “fragmented” group, characterized by small round mitochondria and little branching; 2) a “filamentous” group, with highly connected networks of long/filamentous mitochondria; and 3) an “intermediate” group of cells, containing a mixture of punctate and longer tubular mitochondria. As shown in Figure 4, analysis of the 2D images resulted in quantitative morphological and networking parameters that differed significantly between the three groups. Of note, a more in-depth comparison of the two shape descriptors revealed that FF required smaller sample sizes than AR to detect differences between the three morphological sub-types, and seemed particularly well-suited for distinguishing between cells with filamentous and intermediate mitochondrial morphologies (Figure S4). Likely, this is because AR only measures elongation, whereas FF incorporates the perimeter and therefore is more sensitive to curvature and the irregular shapes of filamentous mitochondria (Figure S4B). Collectively, these results demonstrate that our combined approach for image processing, thresholding, and analysis enables quantitative identification and comparison of mitochondrial morphological sub-types.

**Figure 4:**
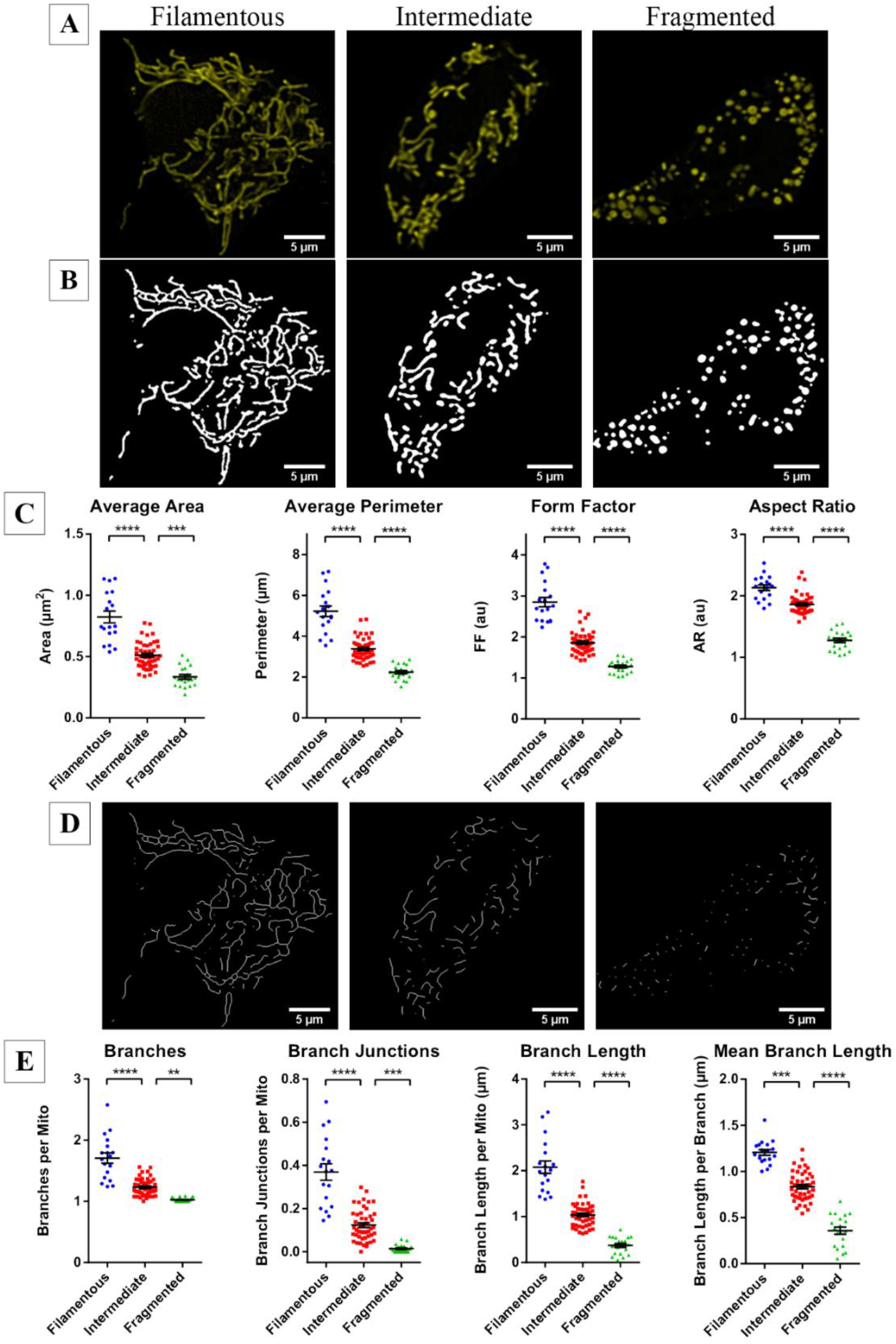
Quantitative comparison of mitochondrial morphology and network connectivity in 2D. Based on visual inspection of their mitochondria, 84 images of Mito-YFP-expressing MIN6 cells were categorized into three morphological groups: fragmented (20 cells), intermediate (46 cells), or filamentous (18 cells). **(A)** Examples of the YFP-labeled mitochondria in representative cells from each group, and **(B)** the objects identified by application of adaptive thresholding to the images. **(C)** The 2D morphological analysis of all cells in each of the categories. **(D)** Skeletonization of the mitochondrial objects identified in panel B. **(E)** Quantitative analysis and comparison of mitochondrial network connectivity performed on all cells in each morphological category. Data are represented by mean ± SEM. One-way ANOVA with Sidak post-hoc test was used to compare the groups; ***p<0.001, ****p<0.0001.

### Validation of morphometric quantifications and classifications by unsupervised clustering

Next, we further tested our pipeline by using Spanning-tree Progression Analysis of Density-normalized Events (SPADE) (32, 33) to obtain an unbiased classification of our test images. The morphological parameters that had been calculated from our image set of 84 mito-YFP-expressing MIN6 cells (shown in Figure 4) were loaded into SPADE, which used these to generate a population tree in which each node represents a cell (Figure 5A). This SPADE tree was then subdivided into 3 cell populations based on automatic classification of their mitochondrial features (Figure 5A; see Materials and Methods for details). When images from each of the three SPADE-identified groups were subsequently examined, the mitochondria in each group were noticeably dissimilar in appearance (Figure 5B), and comparative analysis revealed that there were significant differences in all the morphological descriptors (Figure 5C & D). The morphometric data indicated that SPADE Subgroups 1, 2, and 3 corresponded to cells with filamentous, intermediate, and fragmented mitochondria, respectively. This was confirmed by an 88% match between the unsupervised SPADE clustering and our manual grouping of the cells. Together, these results provide an unbiased validation of the applicability and robustness of our 2D pipeline for analysis of mitochondrial network structure and complexity.

**Figure 5:**
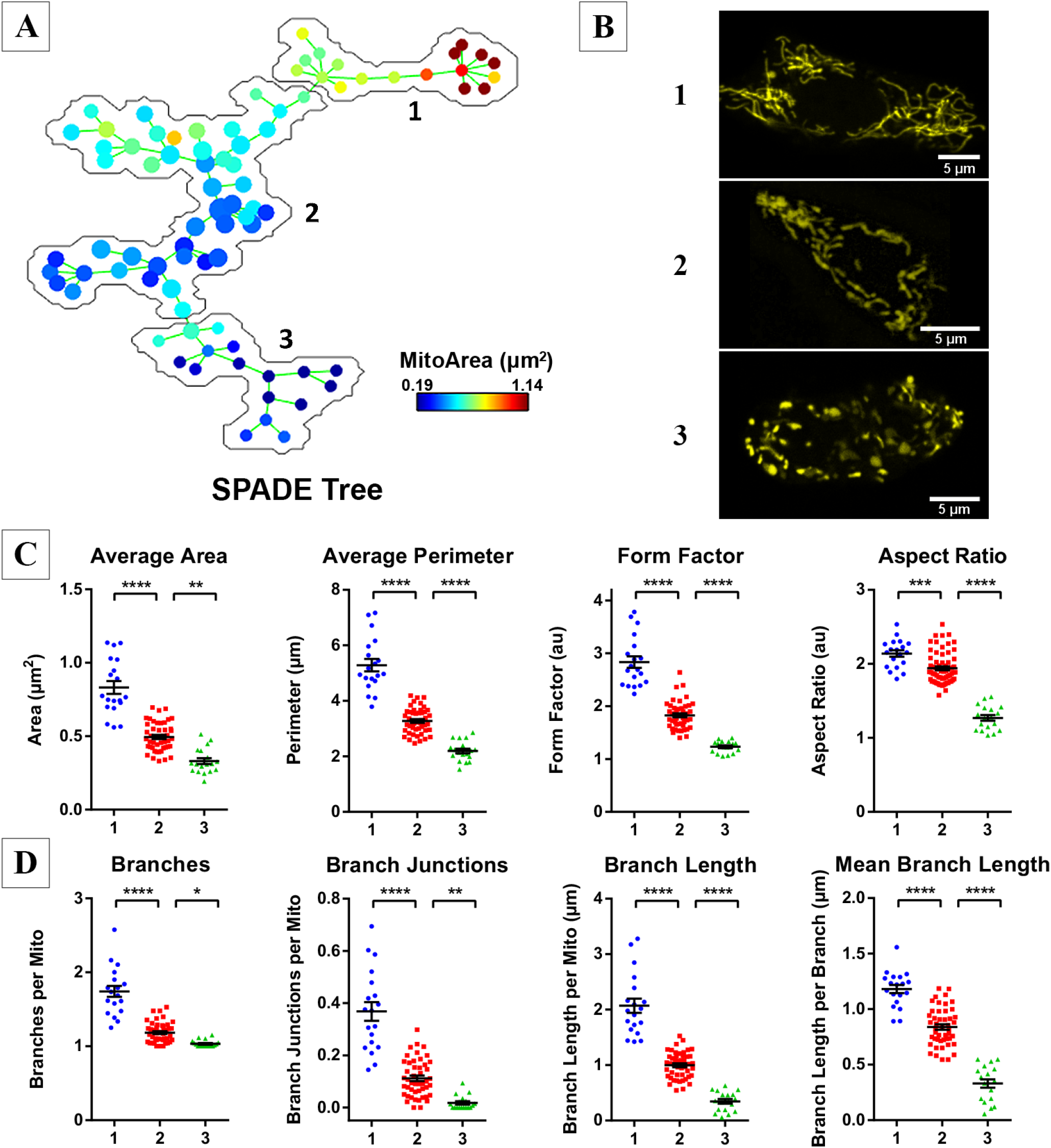
Unsupervised categorization of mitochondrial features using Spanning-tree Progression Analysis of Density-normalized Events (SPADE). **(A)** A SPADE tree was generated based on the same set of 84 images used in Figure 4, and then automatically subdivided into 3 groups; Group 1 contains 19 nodes/cells, Group 2 contains 47 nodes/cells, and Group 3 contains 18 nodes/cells. **(B)** Representative images extracted from each of the three SPADE-generated groups. **(C, D)** Comparison of the mitochondrial morphology and network parameters between the 3 SPADE-identified cell groups. All data are represented by mean ± SEM. *p<0.05, **p<0.01, ***p<0.001, ****p<0.0001 as determined by one-way ANOVA with Sidak post-hoc test; n=84 images.

### Limitations of 2D mitochondrial analysis

Our 2D analyses reliably measure mitochondrial morphology in an optical cross-section and can provide valuable information regarding the state of the organelle. However, when cells are relatively thick and tend to have a mitochondrial network that spans several layers, this approach has its challenges and limitations. It is difficult to know if a given plane in a cell is truly representative, and as illustrated by the green objects in Figure 6A the 2D appearance of a mitochondrion will also depend on its orientation relative to the optical cross-section. Moreover, a 2D image is unlikely to reveal the actual interconnectedness of the mitochondrial network. When a mitochondrion spans multiple planes and intersects the focal cross-section at several points, it can result in a notable misrepresentation of the morphology, as illustrated by the blue schematic object in Figure 6A. That this also occurs in situ is demonstrated in the side-by-side 2D and 3D visualization of a photo-labeled mitochondrion in Figure 6B & C. When viewed in 2D, the localized photo-activation of mito-PAGFP seemed to label four small and distinct mitochondria (Figure 6B; shown in green), but a full 3D reconstruction revealed that it was in fact one continuous organelle (Figure 6C), consistent with diffusion-mediated distribution of GFP within the lumen.

**Figure 6:**
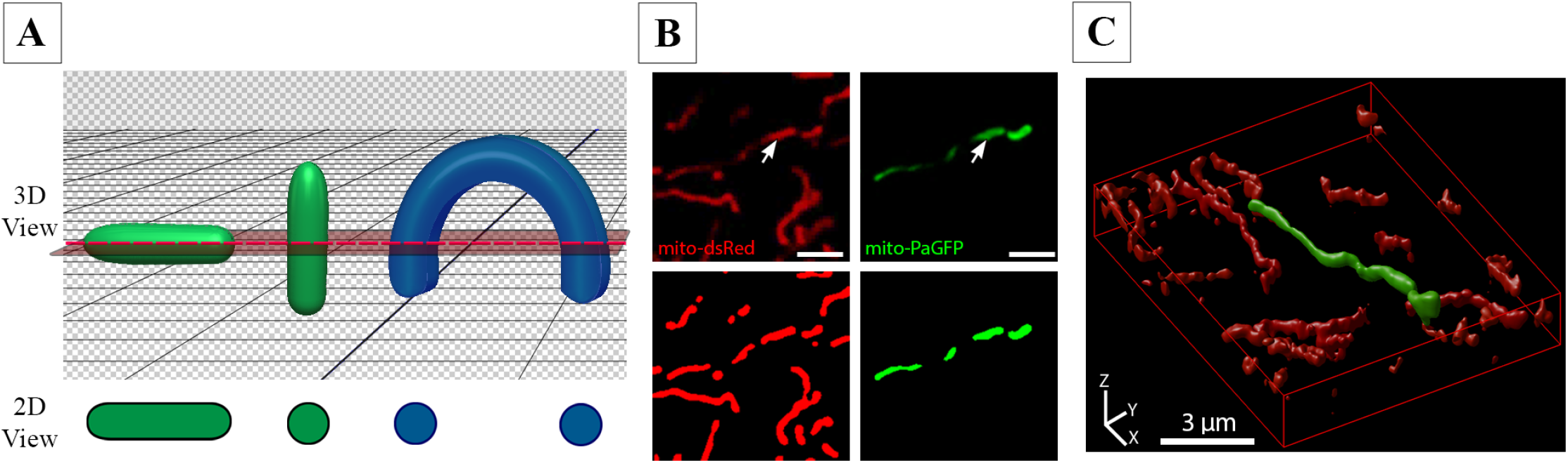
Limitations of 2D morphometric analysis. **(A)** Schematic illustrating the effect of object orientation in 3D space on the image capture in a horizontal 2D slice. The apparent 2D morphology of the same tubular object (shown in green) will depend on its orientation relative to the confocal plane. If a curved object (shown in blue) intersects the confocal plane at several locations, it will erroneously be identified as separate objects. **(B)** MIN6 cells were co-transfected with Mito-dsRed and Mito-PAGFP and photoactivation induced at the point indicated by an arrowhead. Scale bars = 3 µm. *Top row* – 2D image of Mito-dsRed and mito-PAGFP channels after photoactivation. *Bottom row* – objects identified after pre-processing and thresholding of the 2D cross-section. **(C)** Full 3D imaging and reconstruction (rendered using Huygens Professional software) of the same mitochondrial population shown in panel B. Note that the photo-labelled mitochondrion in 2D appears as a series of separate mitochondria, whereas 3D visualization correctly identifies it as one contiguous organelle.

Another inherent limitation of 2D analysis is that it does not allow direct quantitation of the total mitochondrial mass. Cross-sectional area has been used to estimate mass in relatively flat cells like neurons and fibroblasts where mitochondria are confined to a limited number of planes (22, 34). However, this approximation is less appropriate for thicker cells, including β-cells. A common alternative, intended to capture as much of the mitochondrial network as possible, involves acquiring a stack of z-slices and projecting these into a single plane for faster and simpler analysis (6, 23). Such projections contain information from the whole network, but in voluminous cells this will erroneously merge overlapping mitochondria and produce indiscriminate clusters in the resulting image.

As the importance of mitochondrial dynamics and its implication for cellular health and disease has become more apparent, there is also an increasing need for more comprehensive characterization of the organelle. Accordingly, there will inevitably be instances where the caveats of 2D analysis we discussed above become restricting. To enable more precise quantification of mitochondrial volume and network structure we therefore expanded our pipeline to include a complete 3D representation and analysis.

### Three-dimensional imaging and analysis of mitochondria

Full 3D reconstruction of mitochondria can be accomplished by taking a stack of serial slices throughout the volume of the cell and integrating them with software such as ImageJ/Fiji. However, there are technical challenges and constraints specifically associated with 3D imaging. Foremost of these is that the maximum axial resolution (z-axis) of confocal microscopes is approximately 500-800 nm, which is almost three times worse than the lateral (*xy*-plane) (35, 36). As mitochondria are often less than 1 micron in diameter they approach this limit (18). This can lead to a distorted appearance of imaged mitochondria, particularly in the z-axis where it causes artificial stretching and blending of signal from objects in close vertical proximity to each other. In the following section we discuss steps that can be taken to mitigate some of these caveats and improve 3D results.

### Image acquisition and processing requirements for accurate 3D analysis

An important first consideration when acquiring a stack of images for 3D analysis is the z-distance between adjacent imaging planes. If the spacing is too large the final reconstruction will be inaccurate. On the other hand, over-sampling will take unnecessary time, increase photo-toxicity, and require additional resources for image storage and analysis. The distance between serial sections should therefore be set according to the optimal Nyquist sampling rate, which provides the ideal density of information to permit accurate digital reconstruction of an object (37). The Nyquist distance can be calculated using online resources (38).

Even under optimal conditions, a confocal image will be affected by inherent diffraction-induced distortion of the imaged object. This distortion can be represented by a point-spread function (PSF) and then computationally corrected by using deconvolution algorithms. By removing the effects of the PSF, the deconvolution process provides a more correct representation of the underlying object and also helps eliminate out-of-focus light and/or noise in the image (36). In Figure 7 we illustrate this and use the free DeconvolutionLab2 module for ImageJ/Fiji and the commercial deconvolution software, Huygens Professional (SVI), to test the effect of deconvolution on 3D-stacks of mitochondria (see Materials and Methods for details). As seen in Figure 7A, mitochondria in the raw image stack have approximately 2-3x greater diameter in the *xz*-view than in the *xy*-view, which illustrates the z-stretching. The deconvolution algorithms help reduce this distortion, remove noise, and improve the contrast and separation of adjacent objects (Figure 7A and Figure S5). In general, we found that the Huygens deconvolution package reduced axial stretching more effectively than the ImageJ DeconvolutionLab2 module. By and large, however, both deconvolution algorithms significantly increased the quality of 3D mitochondrial network reconstructions compared to the raw confocal images (Figure 7B). Deconvolution also affected subsequent 3D quantifications of mitochondrial number, shape, and size in a way that indicated superior separation of individual mitochondria within the full population (Supplemental Table 1; see discussion of the 3D analysis parameters below). In summary, these results demonstrate that deconvolution of the raw confocal image stacks helps mitigate limitations of 3D imaging and is a necessary step for accurate reconstruction and quantification of the full mitochondrial network.

**Figure 7:**
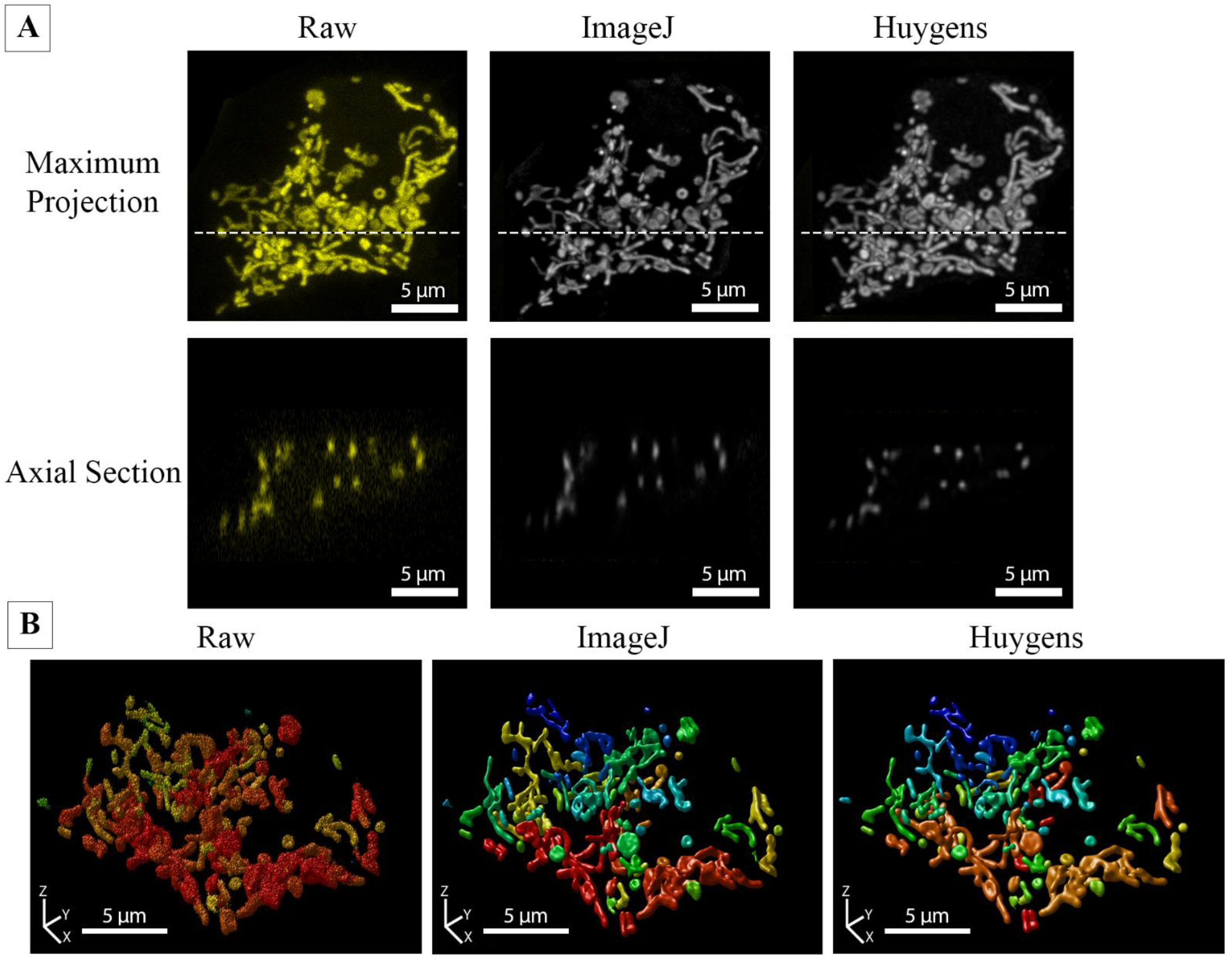
Deconvolution improves the quality and accuracy of 3D mitochondrial analysis. **(A)** A full z-stack was acquired from a Mito-YFP-expressing MIN6 cell that was 11 μm in height. *Top Panel* – Maximum projection views of the z-stack before and after deconvolution. The confocal image stack was deconvolved using either ImageJ DeconvolutionLab (Richardson-Lucy algorithm) or Huygens Professional (Classical Maximum Likelihood Estimation) software for 40 iterations. The dotted line indicates the position of the axial section shown below. *Bottom Panel* – Axial sections (*xz-*plane) of the raw and deconvolved image stacks. The reduction in axial stretching of objects can be seen in the deconvolved stacks, with the best improvement achieved using the Huygens algorithm (see additional details in Figure S5 and Supplemental Table S1). **(B)** 3D renderings of the z-stack before and after deconvolution with ImageJ or Huygens Professional. All 3D visualizations were generated using the Huygens 3D object renderer, with a unique colour assigned to separate objects.

### Three-dimensional quantification of mitochondrial morphology and network connectivity

When a high-quality representation of the full mitochondrial network has been generated, ImageJ/Fiji can be used to extract information about the 3D morphology and connectivity by the same general principles previously discussed for 2D. Mitochondrial size in 3D is represented by volume and surface area, while shape is characterized by the sphericity of the mitochondrial object. The complexity of the 3D network is quantified by the same branch parameters used for 2D (see Figure 3 for a summary). Analogous to our 2D analyses, we evaluated our 3D approach by generating a set of image stacks from mito-YFP-expressing MIN6 cells, and grouping these as fragmented, filamentous, or intermediate based on the visual appearance of the reconstructured mitochondrial networks (Figure 8A). Quantification using ImageJ/Fiji (See Figure 9 and Materials & Methods for details) showed that the number of mitochondria per cell and their average sphericity progressively increased, while the average mitochondrial volume decreased, as we move from filamentous to intermediate to fragmented morphologies (Figure 8B). In contrast, the total mitochondrial volume of each cell remained constant, highlighting that significant morphological heterogeneity can occur independent of changes to mitochondrial mass (Figure 8B). In the skeletonized network the number of branches and branch junctions progressively decreased, illustrating that mitochondrial fragmentation, not surprisingly, is associated with a reduction in overall network complexity (Figure 8C & D). Together, the above results and discussions demonstrate how standard confocal imaging can be combined with ImageJ/Fiji-based processing and analysis, to quantify volume, morphology, and connectivity of the entire mitochondrial network in pancreatic β-cells. To our knowledge, full 3D characterization of live β-cell mitochondria has previously only been done at this level using specialized super-resolution imaging techniques (18, 24).

**Figure 8:**
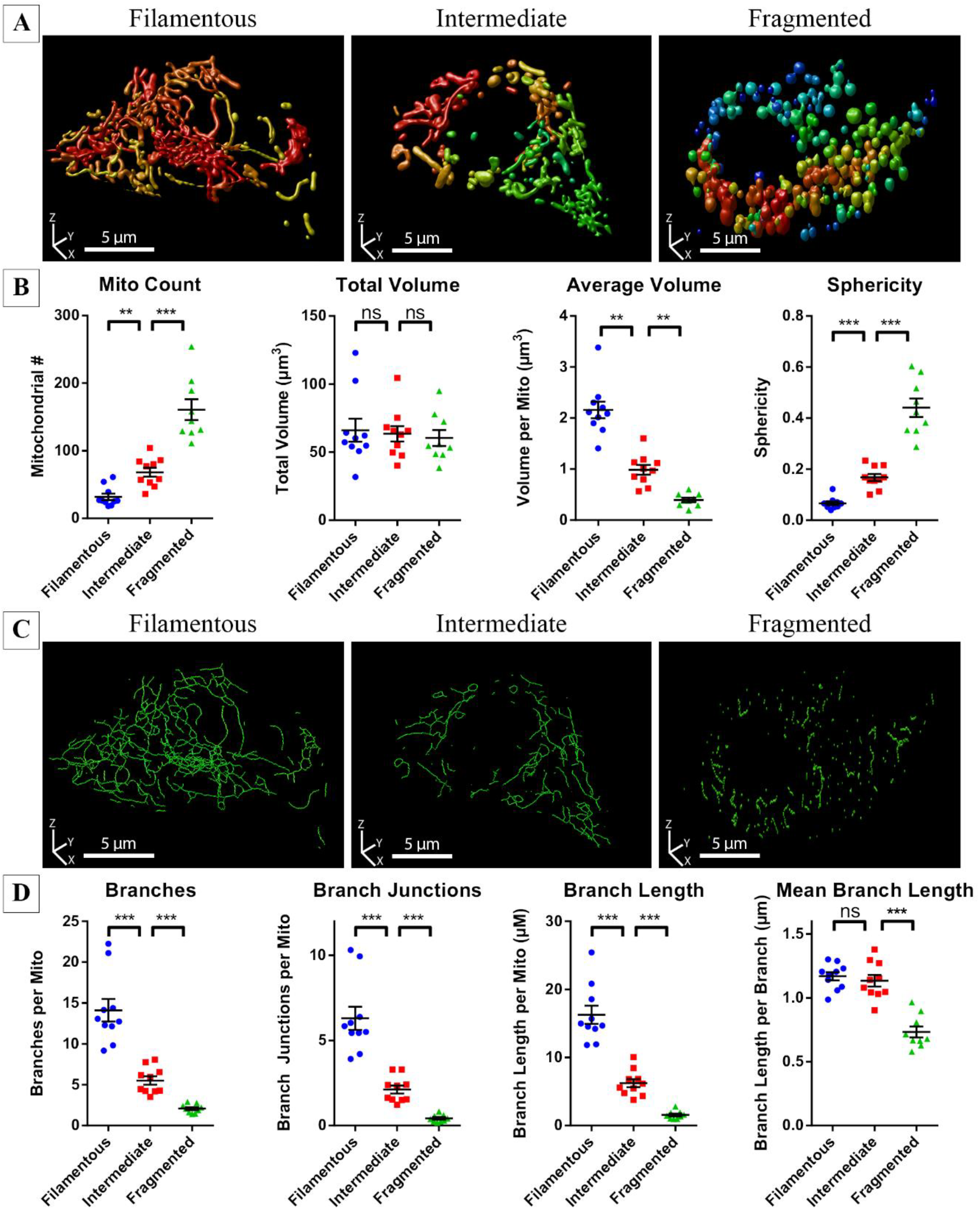
Quantitative comparison of mitochondrial morphology and network connectivity in 3D. Image stacks of Mito-YFP-expressing MIN6 cells were visualized in 3D and their mitochondria manually categorized as fragmented, intermediate or filamentous. **(A)** 3D renderings (produced using Huygens Professional) of representative Mito-YFP-expressing MIN6 cells from each of the morphological categories. **(B)** Quantitative 3D analysis and comparison of mitochondrial morphology between cells in each category. **(C)** 3D renderings of the skeletonized mitochondrial network of the cells depicted in panel A. **(D)** Quantitative 3D analysis and comparison of mitochondrial network connectivity between cells in each category. All data are represented by mean ± SEM. *p<0.05, **p<0.01, ***p<0.001, as determined by one-way ANOVA with Sidak post-hoc test. N=10 cells in each category.

**Figure 9:**
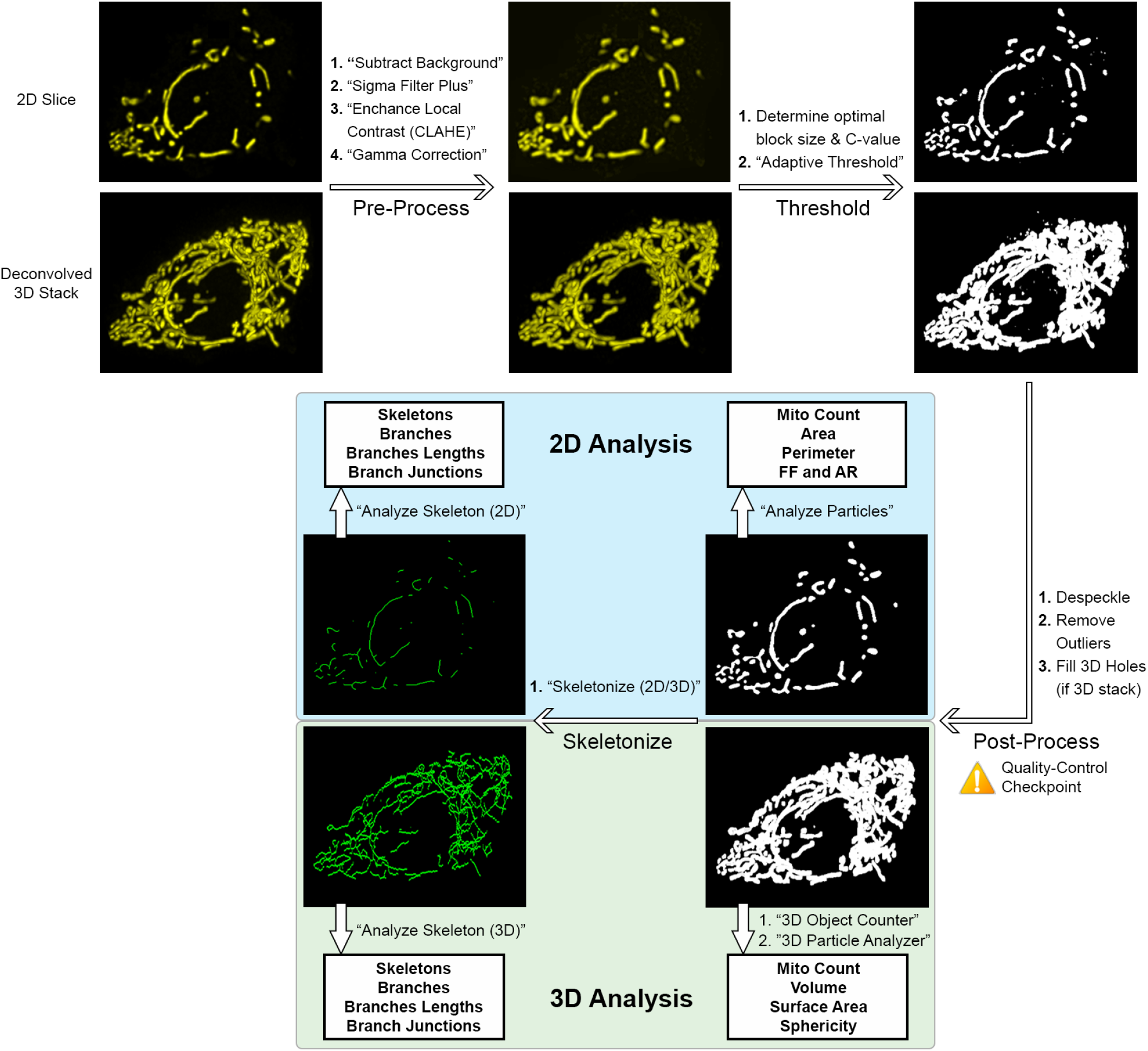
Summary of pipeline for 2D and 3D mitochondrial analysis in ImageJ/Fiji. For illustration, an image stack was acquired from a MIN6 cell expressing Mito-YFP; a representative slice is shown as the 2D input, and the entire stack (after deconvolution) as the 3D input. 3D Stacks are represented as maximum projections here. Scale bars = 5 μm. See main text and Materials & Methods for additional details and parameter values.

### Pipeline Summary

Figure 9 illustrates the overall pipeline for 2D and 3D mitochondrial analysis. In summary, 2D image slices or 3D image stacks are first acquired, and the latter deconvolved prior to analysis. In ImageJ/Fiji, deconvolution of 3D stacks is done using the DeconvolutionLab2 module (39) and if desired, the 3D stack can be visualized using the “3D Viewer” or “Volume Viewer” functions. Alternatively, 3D deconvolution and visualization can be done using commercial software, such as Huygens, if available to the user (Figure 7). For analysis, all images are then pre-processed using the commands: “Subtract Background”, “Sigma Filter Plus”, “Enhance Local Contrast”, and “Gamma Correction”. We then empirically test a range of block sizes and C-values for the “Adaptive Threshold” command to establish the optimal values and use these as input when applying the threshold algorithm. The resulting binarized images are post-processed using the “Despeckle”, “Remove Outliers”, and “Fill 3D Holes” commands. At this stage, we recommend comparing the final thresholded image to the original images as a quality control check of the object identification and segmentation. The identified mitochondrial objects are then analyzed in 2D using “Analyze Particles”, which provides mitochondrial count, area, perimeter, form factor (FF), and aspect ratio (AR). For 3D analysis, we use the “3D Object Counter” and “3D Particle Analyzer” (from the MorphoLibJ package) commands to quantify count, volume, surface area, and sphericity. The thresholded objects are then converted into skeletons using “Skeletonize (2D/3D)”, and we apply the “Analyze Skeleton” command to obtain the number of skeletons, number of branches, length of branches, and number of branch junctions in the 2D or 3D network. Additional details and parameter values can be found in Materials and Methods.

### Quantifying physiological and pathophysiological changes to mitochondrial morphology and networking

Having established and validated the mitochondrial analysis pipeline, we next used it to characterize mitochondrial changes under relevant physiological and pathophysiological conditions. As a test of acute functional responses, primary mouse β-cells were cultured in either basal (3 mM) or stimulatory (17 mM) glucose for 1 hour and co-stained with MitoTracker green (MTG) and the mitochondrial membrane potential-sensitive dye TMRE (Figure 10A). The MTG fluorescence is insensitive to changes in mitochondrial polarization and served as the signal for mitochondrial detection and morphological characterization (40). The TMRE intensity provided a simultaneous readout of the activity of the individual mitochondrial units, and as expected stimulatory glucose increases the TMRE/MTG intensity ratio (Figure 10B). By visual inspection there were no obvious differences in mitochondrial morphology between the cells in low and high glucose (Figure 10A), but quantitative analysis revealed a number of significant effects (Figure 10C & D). Despite no change to total mitochondrial area, glucose stimulation increased the number of mitochondria, reduced their average size (area and perimeter) and made them more round (decreased form factor); all of which suggests increased mitochondrial fission (Figure 10C). This was further supported by skeletonization analysis, which showed that stimulatory glucose caused an overall reduction in mitochondrial network connectivity (decreased branch parameters) (Figure 10D). This experiment agrees with previous reports linking drp1-dependent mitochondrial fission to glucose-stimulated insulin secretion (19, 20), and demonstrates that our analysis pipeline is sensitive enough to allow quantitative detection of subtle physiological changes to mitochondrial morphology and networking.

**Figure 10:**
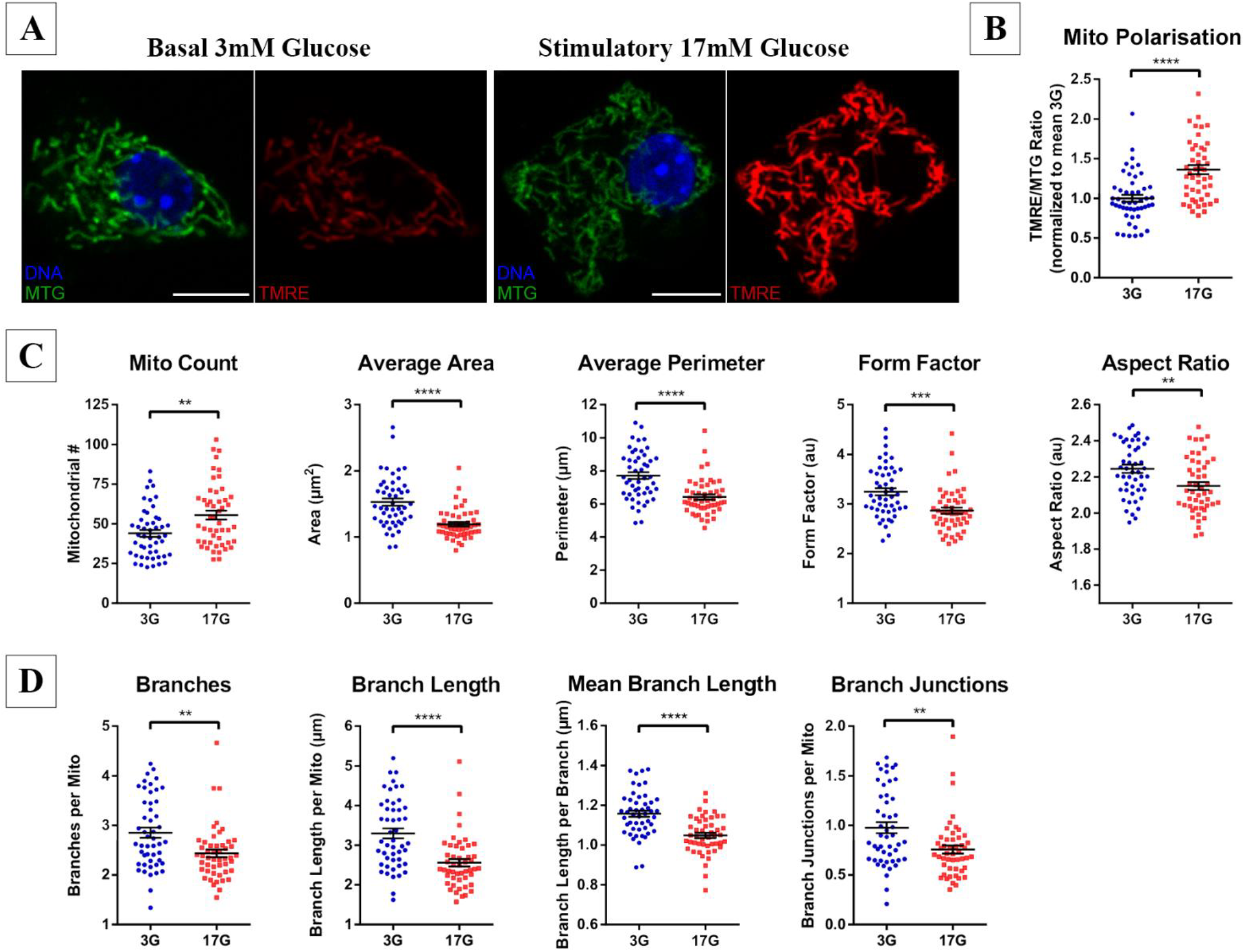
Two-dimensional analysis shows that glucose stimulation is associated with mitochondrial fission in pancreatic islet cells. Dispersed mouse islet cells were treated in either 3mM glucose (3G) or 17mM glucose (17G) for 60 minutes and then labeled with Hoescht 33342, MitoTracker Green FM (MTG), and TMRE before 2D imaging. **(A)** Representative images of a MTG and TMRE stained islet cell in 3G and 17G. **(B)** TMRE/MTG ratio (normalized to average 3G), indicating the degree of mitochondrial hyperpolarisation. Mitochondrial morphology and polarization were quantified using our 2D analysis pipeline in Fiji/ImageJ (see Materials and Methods). Comparison of mitochondrial morphometry **(C)**, and mitochondrial network connectivity **(D)** demonstrates significant differences between cells acutely treated with low and stimulatory glucose. All data are represented by mean ± SEM. **p<0.01, ***p<0.001, ****p<0.0001 as determined by Student’s t-test; n=49 cells in each glucose treatment from 4 mice.

As an example of a full 3D application, we quantified the mitochondrial changes in palmitate-treated MIN6 cells; an *in vitro* model of the β-cell lipotoxicity associated with obesity and type 2 diabetes. As expected from previous 2D analyses (6) we observed a fragmentation of the mitochondrial network following treatment with a high concentration of palmitate (Figure S6). This pathophysiological stress response did not affect total mitochondrial volume but was clearly reflected in all parameters describing the 3D shape and size of individual mitochondrial units (Figure S6). Comparing the 2D morphological changes associated with 1 hour of glucose stimulation and 6 hours of palmitate exposure, it is interesting to note that palmitate treatment reduced AR by 26% and FF by 29%, while glucose stimulation only decreased AR and FF by 4% and 12%, respectively. This suggests that physiological fission generates daughter mitochondria that largely retain their shape, in contrast to the more pronounced stress-induced fragmentation, which also causes a striking rounding of the smaller organelles.

### Four-dimensional analysis of mitochondrial dynamics

At any given time, the overall structure of a mitochondrial network reflects the net balance of fusion and fission between individual mitochondria. These are dynamic, energy-dependent, processes that involve mitochondrial movement, and coordinated actions of proteins that mediate fusion of the outer and inner membranes, or constriction and splitting of the organelle (2). Based on static image analysis alone it can be difficult to know the reason for a change in morphometry. For instance, a more connected and elongated network can be the result of an increase in fusion events, a decrease in fission activity, or a combination of both. To further understand the underlying changes, it can therefore be valuable to monitor mitochondrial movement, morphological changes, and organelle interactions in real-time. In practical terms, this requires that image acquisition can be repeated at sufficiently frequent intervals, and that the analysis is extended to the time-domain. Previous studies have applied these principles to 2D images to provide important insights regarding mitochondrial dynamics and turnover in pancreatic β-cells (3, 6).

Here, we tested the feasibility of recording and quantifying the time-dependent dynamics of the full 3D mitochondrial network (i.e. an extension to 4D analysis). For this, we expressed mito-YFP in MIN6 cells and imaged these in a stage-top incubator on the confocal microscope. 3D time-lapse data were generated by acquiring z-stacks of the cells at regular time-intervals (every 45 s) for a period of 30 minutes. At the 13-minute mark, we added a high concentration of the mitochondrial uncoupling agent, FCCP, with the purpose of inducing a relatively rapid change in mitochondrial dynamics and architecture. As seen in Figure 11 and Supplemental Video 1, the FCCP triggered a rapid, and dramatic, loss of mitochondrial connectivity along with an increase in the number of organelles. Interestingly, there was also a transient decrease in both average and total mitochondrial volume, which indicates an initial contraction and shrinking of the mitochondrial fragments followed by significant swelling; a known response to stress and osmotic shock (41). The abrupt and severe deterioration of the mitochondrial network likely reflects the induction of apoptosis due to profound damage from high levels of FCCP.

**Figure 11:**
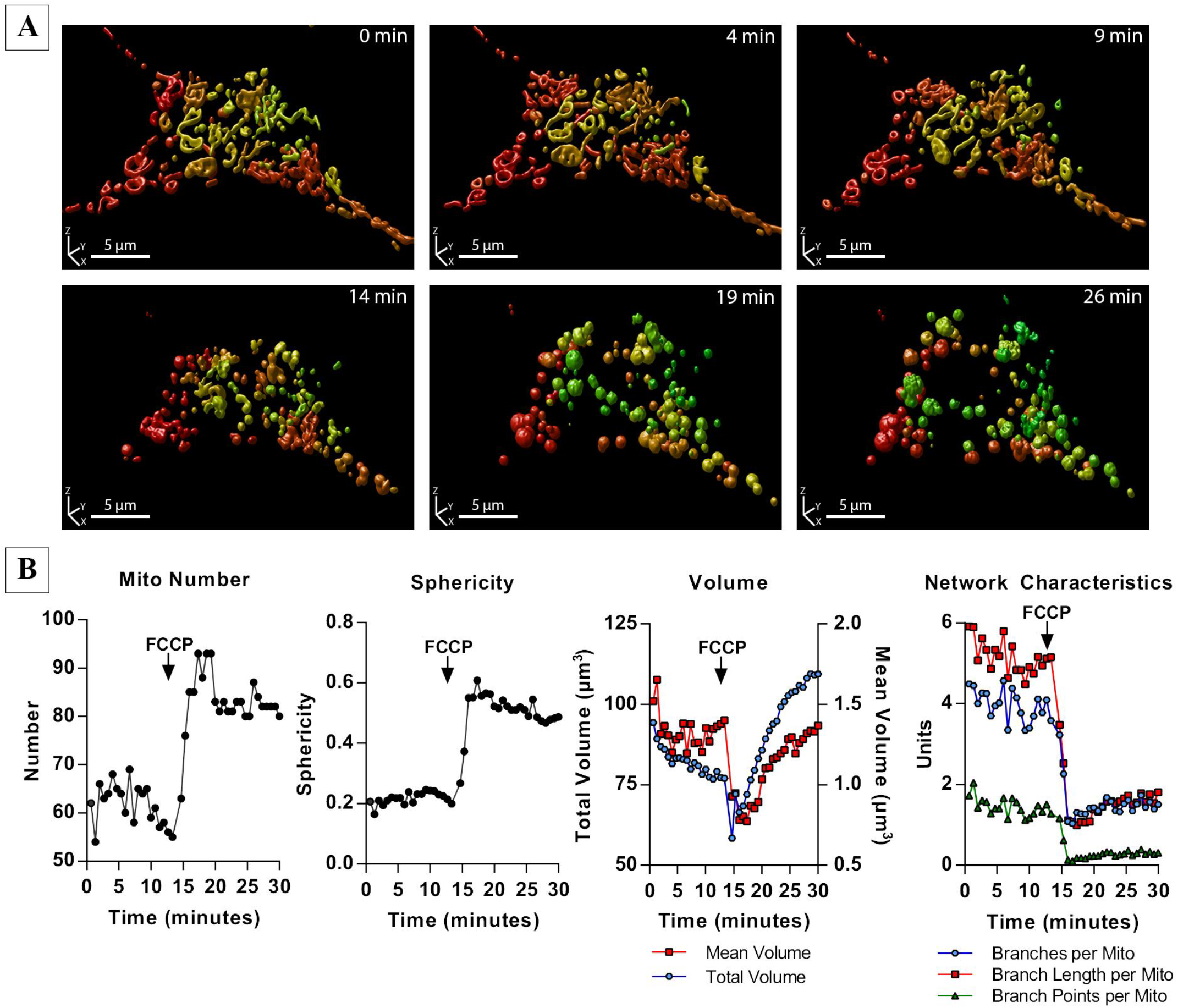
Time-lapse 3D imaging (*xyzt*) and analysis of mitochondrial dynamics. Mito-YFP-expressing MIN6 cells were imaged in a stage-top incubator with one full image stack acquired every minute. **(A)** 3D renderings (produced in Huygens Professional) of the mitochondrial network in a single cell at different time-points. A high concentration FCCP (25 µM) was added to the incubation media around the 13-minute mark. **(B)** Quantitative analysis of the time-dependent effects of FCCP on mitochondrial number, sphericity, total and mean volume, and network characteristics in the cell.

With this proof-of-principle experiment we have established the feasibility of analyzing the temporal dynamics of a full mitochondrial network using standard confocal microscopy. A powerful next step could be to combine this 4D approach with other tools such as photo-labeling and tracking of individual organelles, to generate even more complete and in-depth knowledge of the events that shape the mitochondrial network in health and disease.

## CONCLUSION & PERSPECTIVES

Most aspects of cellular function and survival are linked to mitochondrial physiology or signals originating from the organelle, and in these contexts the importance of mitochondrial morphology and dynamics has become evident (1, 2). The integrity of the organelle itself, and by extension the metabolic health of the cell, depends on the capacity for mitochondrial adaptation to stress and on selective turnover of damaged parts of the network by mitophagy (3). These processes rely on mitochondrial fusion and fission dynamics, which require sensitive live-cell imaging approaches to study (42). Our current understanding of mitochondrial dynamics in pancreatic β-cells has also been based largely on such imaging approaches (3, 6). However, it is challenging to accurately quantify β-cell mitochondrial morphometry and dynamics by fluorescence microscopy, and many important questions remain unanswered.

In the previous sections, we established a comprehensive set of methods for quantitative image analysis and ‘morphofunctional’ characterization of mitochondria based on standard confocal microscopy and the ImageJ/Fiji shareware. The robustness of these approaches was validated in several ways, including by unsupervised data clustering. We demonstrated the applicability of the resulting pipeline for cells with dense multi-layered mitochondria by conducting detailed 2D and 3D morphometric analyses of β-cells, and further extended these to 4D time-lapse imaging with the accuracy needed for quantitative assessment of network dynamics. To help researchers implement these methods, we have also built our analysis pipeline into a plugin for ImageJ/Fiji called Mitochondria Analyzer. The plugin is publicly available (28) and includes a graphical user interface to facilitate pre-processing, parameter optimization, image thresholding and automated morphofunctional analysis of mitochondrial images or image stacks, according to the work-flow we have presented (Figure 9).

When testing the pipeline, we demonstrated the capability for multi-parameter characterization by performing 2D analyses of β-cells co-stained with MTG and TMRE for simultaneous recordings of changes to mitochondrial morphology and membrane potential. However, the pipeline can in principle be applied to any number of mitochondrial parameters, provided they can be jointly imaged and then quantified using shape- and intensity-based descriptors. We therefore predict that the same type of analysis using a stable mitochondrial label combined with one or more spectrally distinct fluorescent biosensors, e.g. for mitochondrial redox state, matrix Ca^2+^ or pH, could provide valuable insights into physiological and pathophysiological structure-function relationships in mitochondria. Importantly, the analysis pipeline treats all identified objects separately, and can therefore extract the morphological and functional descriptors on a per-mitochondria basis. In the previous sections we presented our results based on the cellular averages, but the same data-sets contain the descriptors associated with thousands of individual mitochondria and can be mined for a wealth of information about morphometry-physiology correlations and heterogeneity at the organelle level (43, 44). It should also be emphasized that the practical considerations and best-practices we have discussed, and incorporated into our pipeline, are not restricted to β-cell analyses, but can also be applied to other cell types.

Finally, the pipeline can in principle also be used to investigate fluorescently labelled organelles other than mitochondria, provided that appropriate thresholding/analysis parameter adjustments can be made. The importance of inter-organelle contacts for cellular function and health are becoming clear, as is the highly complex and dynamic nature of the “organelle interactome” (45–47). Within the technical boundaries associated with standard confocal microscopy, the analysis approaches we have described here can help most research laboratories achieve the level of accuracy needed to explore internal mitochondrial network interactions, and likely also the mitochondrial relationship with other organelles, as we work to clarify the sub-cellular basis of diabetes and other diseases.

## MATERIALS AND METHODS

### Reagents

Collagenase type XI (#C7657), Tetramethylrhodamine Ethyl Ester (TMRE, #87917), D-glucose (#G7528), Bovine Serum Albumin (BSA, #A7030), FCCP (#C2920) and palmitic acid (#P5585) were purchased from Sigma-Aldrich (St. Louis, Missouri). MitoTracker Deep Red FM (#M224726), MitoTracker Green FM (MTG, #M7514), Hoechst 33342 (#H3570), RPMI 1640 (#11879), Dulbecco’s Modified Eagle’s Medium (DMEM, #11995), Fetal Bovine Serum (FBS, #10438), Trypsin-EDTA (#25300), Penicillin-Streptomycin 10,000 U/mL (#15140), and HBSS (#14185) were purchased from Life Technologies/Thermo Fisher Scientific (Carlsbad, California). Dimethyl sulfoxide (DMSO, #BP231) was purchased from Fisher Scientific (Waltham, Massachusetts). Minimum Essential Media (MEM, #15-015-CV) was purchased from Corning (Corning, New York). The mitochondria-targeted YFP (mito-YFP) and mitochondria-targeted photoactivatable GFP (mito-PAGFP) plasmids were gifts from Dr. Mark Cookson (48) and Dr. Richard Youle (Addgene #23348) (5), respectively.

### Cell Isolation and Culture

MIN6 cells were cultured at 37°C and 5% CO_2_ using complete DMEM supplemented with 10% FBS and 2% Penicillin-Streptomycin. Culture media was replaced every 2 days and cells were passaged upon reaching 70-80% confluency. To assess the effects of palmitate on mitochondrial networks, cells were cultured for 6 hours in complete DMEM supplemented with either 1.5 mM palmitate complexed to BSA in a 6:1 ratio, or BSA-only vehicle control.

Pancreatic islets were isolated from wild-type male mice of a mixed C57BL/6 and CD1 background using collagenase digestion and filtration-based purification, as previously described (49). The isolated islets were hand-picked and allowed to recover overnight before being dispersed into single cells and seeded on 25 mm glass coverslips (50). The islet cells were cultured in RPMI completed with 10% FBS and 2% Penicillin-Streptomycin at 37ºC and 5% CO_2_ for 4 days before imaging. All animal procedures were approved by the University of British Columbia Animal Care Committee.

### Cell Transfection and Mitochondrial Labeling

MIN6 cells were seeded at a density of 2.0 × 10^5^ on 25 mm glass coverslips (0.13-0.16 mm thickness, VWR #16004-310) and incubated for 24 hours before being transfected with mito-YFP, mito-dsRed, or mito-PAGFP plasmids using Lipofectamine® 2000 (Life Technologies #11668), as per the manufacturer’s protocol. All plasmids were expressed for at least 24 hours before confocal microscopy.

To assess the effect of acute glucose exposure on mitochondrial morphology and membrane potential, primary mouse islet cells were cultured for 60 minutes in completed RPMI media containing 3 mM glucose or 17 mM glucose and then stained with 0.1 µg/mL Hoescht 33342, 50 nM MTG, and 25 nM TMRE for 30 min, followed by a wash with completed RPMI immediately prior to imaging.

### Image Acquisition by Confocal Microscopy

Live cells were imaged in a Tokai Hit INUBTFP-WSKM stage-top incubator at 37°C on a Leica SP8 Laser Scanning Confocal Microscope (Concord, Ontario, Canada). For 2D, images were acquired using a 63x HC Plan Apochromatic water immersion objective (1.2 NA). Pixel size was adjusted using the “Optimize” function in the Leica LASX Software and the pinhole size was 1.0 AU. For 3D acquisition, *z*-stacks were obtained using a 63x oil immersion objective (1.4 NA). Pixel size (*x*, *y*) and *z-*spacing were adjusted as per the calculated optimal Nyquist sampling parameters (38), and pinhole size was reduced to 0.75 AU. The *z*-step size generally varied between 170-220 nm. Bi-directional scanning was enabled, and all images were acquired using at least three frame averages. Laser power, detector filtering/gating, and gain were adjusted to maximize signal without saturation, while also minimizing background signal, cross-fluorescence, and photobleaching.

### Time-Lapse 3D Imaging

Time-lapse 3D (*xyzt*) imaging was performed on MIN6 cells transfected with mito-YFP plasmid. Acquisition settings were established as above, and *z*-stacks were acquired every 45 s for 30 min. The acquisition time for each stack was approximately 30 s. At the 13 min-mark, FCCP was added to the chamber for a final concentration of 25 µM.

### Mitochondrial labeling by Photoactivatable Green Fluorescent Protein

MIN6 cells were co-transfected with mito-PAGFP and mito-dsRed 24 hours prior to imaging, as described above. Mito-dsRed was visualized using a 561 nm excitation laser with emission detected between 585 nm to 650 nm. Individual mitochondria were marked for photo-labeling using the “Bleach Point” function in the Leica LasX software, and the PAGFP was activated using a 405 nm laser pulse of 150 ms duration. The activated PAGFP was then imaged using a 488 nm excitation laser with an emission range between 505 nm and 550 nm. All other imaging settings were as described above.

### Image Deconvolution

Deconvolution of 3D and 4D stacks was performed in Huygens Professional version 16.10 (SVI) using the CMLE (Classic Maximum Likelihood Estimation) algorithm with a Signal-to-Noise Ratio of 7.0, maximum iterations of 40, and quality threshold of 0.001. For deconvolution in ImageJ/Fiji, the “PSF Generator” plugin (51) was used to generate a theoretical PSF based on our microscope parameters, and deconvolution was performed with the “DeconvolutionLab2” module utilizing the Richardson-Lucy TV algorithm with regularization set to 0.0001 and maximum iterations of 30 (39, 52).

### Image Processing and Thresholding

The workflow and procedures for image processing and thresholding are summarized in Figure 9. Using ImageJ/Fiji, 2D images or deconvolved 3D image stacks (operating on each slice in the stack) were pre-processed using the following commands: 1) “Subtract Background” (radius = 1μm) to remove background noise; 2) “Sigma Filter Plus” (radius = 0.1 μm, 2.0 sigma) to reduce noise and smooth object signal while preserving edges; 3) “Enhance Local Contrast” (block size = 64, slope = 2.0 for 2D and 1.25 for 3D stacks) to enhance dim areas while minimizing noise amplification; and 4) “Gamma Correction” (value = 0.80 for 2D and 0.90 for 3D) (29) to correct any remaining dim areas. To identify mitochondria in the images, we evaluated multiple global and local thresholding algorithms (Figure 2 and Figures S1-S3). Based on our comparisons, we elected to use the “Adaptive Threshold” method (30). In the “Adaptive Threshold” plugin, block size was set to an equivalent of 1.25 μm and the optimal C-value was empirically determined for each image set (See Figure S2 for additional details). The thresholded images were then post-processed using “Despeckle” and then “Remove Outliers” (radius = 0.15 μm^2^) to remove residual noise. For 3D stacks we additionally applied the “Fill 3D Holes” command from the “3D ROI Manager” plugin (53).

### 2D Analysis of Mitochondrial Function, Morphology, and Network Characteristics

The approach for quantification of mitochondrial characteristics is summarized in Figure 9. For 2D analysis, the image was first processed and thresholded (see above) and the resulting binary image was used as the input for the “Analyze Particles” command (Size = 0.06 μm^2^-Infinity, Circularity = 0.00-1.00), measuring for “Area”, “Perimeter”, and “Shape Descriptors”. Form Factor (FF) was derived as the inverse of the “Circularity” output value. For network connectivity analysis, the “Skeletonize 2D/3D” command was applied to the thresholded image to produce a skeleton map, and the “Analyze Skeleton” command was used to calculate the number of branches, branch lengths, and branch junctions in the skeletonized network.

To simultaneously measure mitochondrial polarisation and morphology, islet cells were co-stained with MTG and TMRE. Our threshold method was first applied to the MTG channel and morphological analysis was done on the identified objects. Additionally, the “Analyze Particles” command (“Add to Manager” option enabled) was used to convert the identified objects (mitochondria) into regions of interest (ROIs). These ROIs were then superimposed onto the raw images of the MTG and TMRE channels, and the MTG and TMRE intensities of each individual mitochondrion were measured as the “Mean gray value” obtained via the “Analyze Particles” command. The degree of mitochondrion polarisation was then expressed as the ratio of TMRE to MTG intensity and correlated with mitochondrial morphology on a per-organelle basis.

### 3D Analysis of Mitochondrial Morphology and Network Characteristics

For 3D analysis, the image stacks were first deconvolved, pre-processed, and thresholded as described above and summarized in Figure 9. Next, the “3D Object Counter” command (Size = 0.6 μm^3^-Infinity) was used to calculate the number of mitochondrial objects and produce a labelled object map. The object map was subsequently used as an input for the “Particle Analyzer 3D” command (part of the MorphoLibJ package) (54) to calculate the volume, sphericity (weighted by volume of the object), and corrected surface area (“Crofton 13 directions” method) of each mitochondrial object. Network connectivity analysis was performed on the skeletonized 3D network using the same commands as 2D analysis. For 4D (*xyzt*) analysis, these 3D analysis steps were performed on each stack obtained in the time-course acquisition.

### Unsupervised Categorization of Mitochondrial Morphology using SPADE

Spanning-tree Progression Analysis of Density-normalized Events (SPADE v3.0) (32, 33) was used to automatically classify 2D mitochondrial images into 3 different categories based exclusively on their calculated morphological and network parameters. Briefly, mitochondrial parameter data was transformed into a Flow Cytometry Standard (FCS) file using FlowJo V10 and loaded into SPADE. A SPADE tree was created using default settings, without application of arcsinh transformation or removal of outliers. The number of desired clusters was set equal to the total number of images in the data set. The “Auto Suggest Annotation” function was then used to partition the SPADE tree into 2 subgroups and the larger of these was subsequently auto-partitioned again, resulting in a total of 3 subgroups. The data in these SPADE-identified groups were then exported as CSV formatted files for statistical comparison.

### Statistical analysis

All data were represented as mean ± standard error of the mean (SEM). Data were analyzed in GraphPad Prism 6.0 software (La Jolla, California) using Student’s t-test or One-way ANOVA, followed by Sidak multiple comparison test as appropriate. Statistical significance was set at a threshold of p < 0.05.

## ACKNOWLEDGEMENTS

We thank Dr. Jingsong Wang, Mei Tang, and Mitsuhiro Komba at BC Children’s Hospital Research Institute (BCCHRI) for technical assistance and Daniel J. Pasula for constructive feedback and discussion. This work was supported by operating funds from the Canadian Institutes of Health Research (CIHR; MOP-119537) and JDRF (2-2013-50). Research infrastructure was funded by the Canadian Foundation for Innovation (CFI; #30214). DSL received salary support from BCCHRI and Diabetes Canada. RS was supported by a Canada Graduate Scholarship from CIHR.

## AUTHOR CONTRIBUTIONS

AC, RS and DSL all contributed to the design of methods and validations. AC and RS performed the experiments and validations. AC, RS and DSL all analyzed/interpreted the data and wrote the paper. AC developed the associated Mitochondria Analyzer plugin for ImageJ.

