## Supplemental Figures S1-S6 and Table S1 for "A pipeline for multidimensional confocal analysis of mitochondrial morphology, function and dynamics in pancreatic β-cells"

### Chaudhry et al. – Supplemental Figure 1

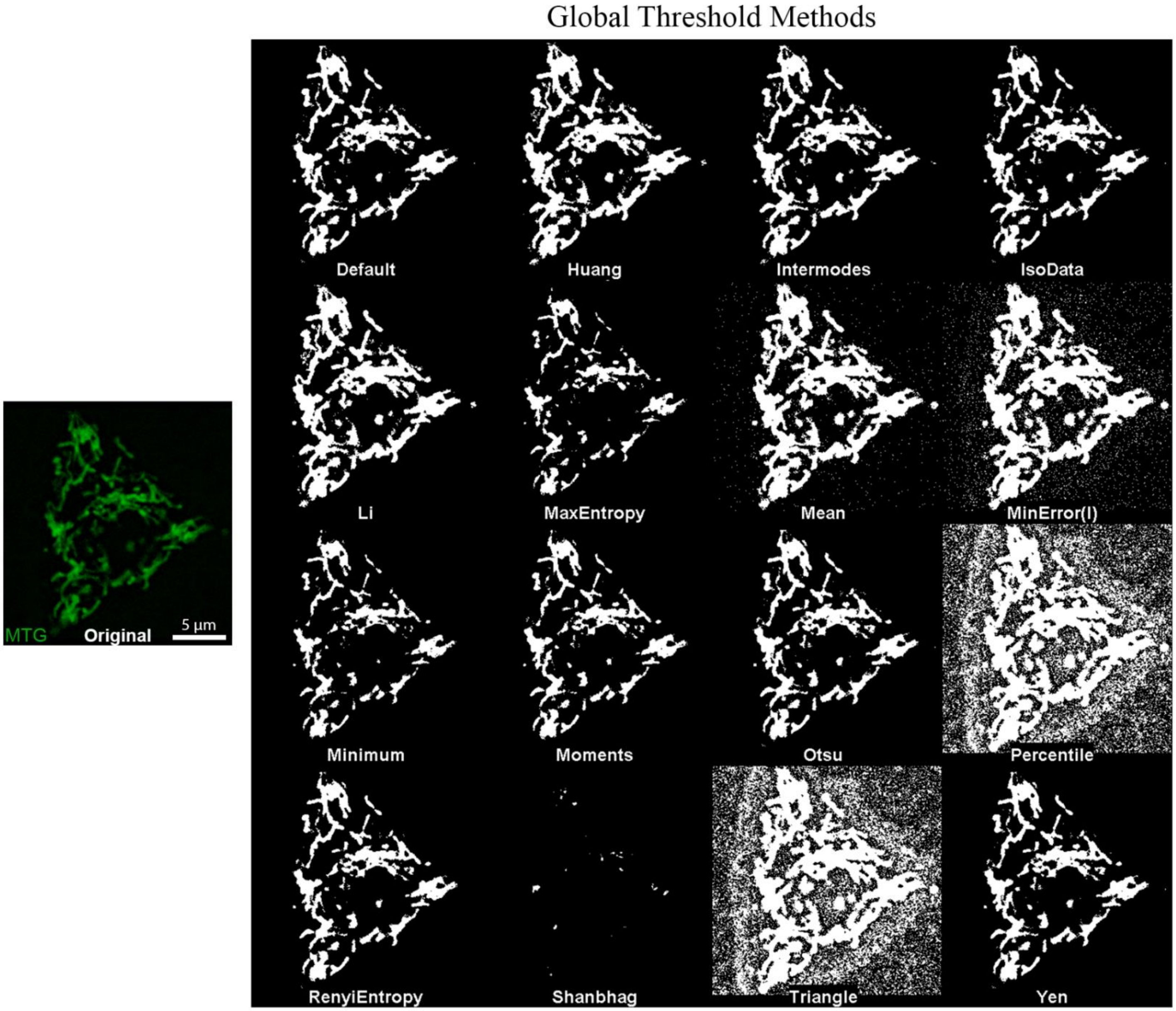


**Figure S1: Comparison of the global thresholding algorithms available in ImageJ/Fiji.** Global algorithms identify a threshold value based on the intensity histogram of the full image. *Left* - The original 2D image of an islet cell stained with MitoTracker Green (MTG). *Right* – Montage showing the results of thresholding the original image using the different global thresholding methods available under the ImageJ/Fiji 'AutoThreshold' command.

### Chaudhry et al. – Supplemental Figure 2

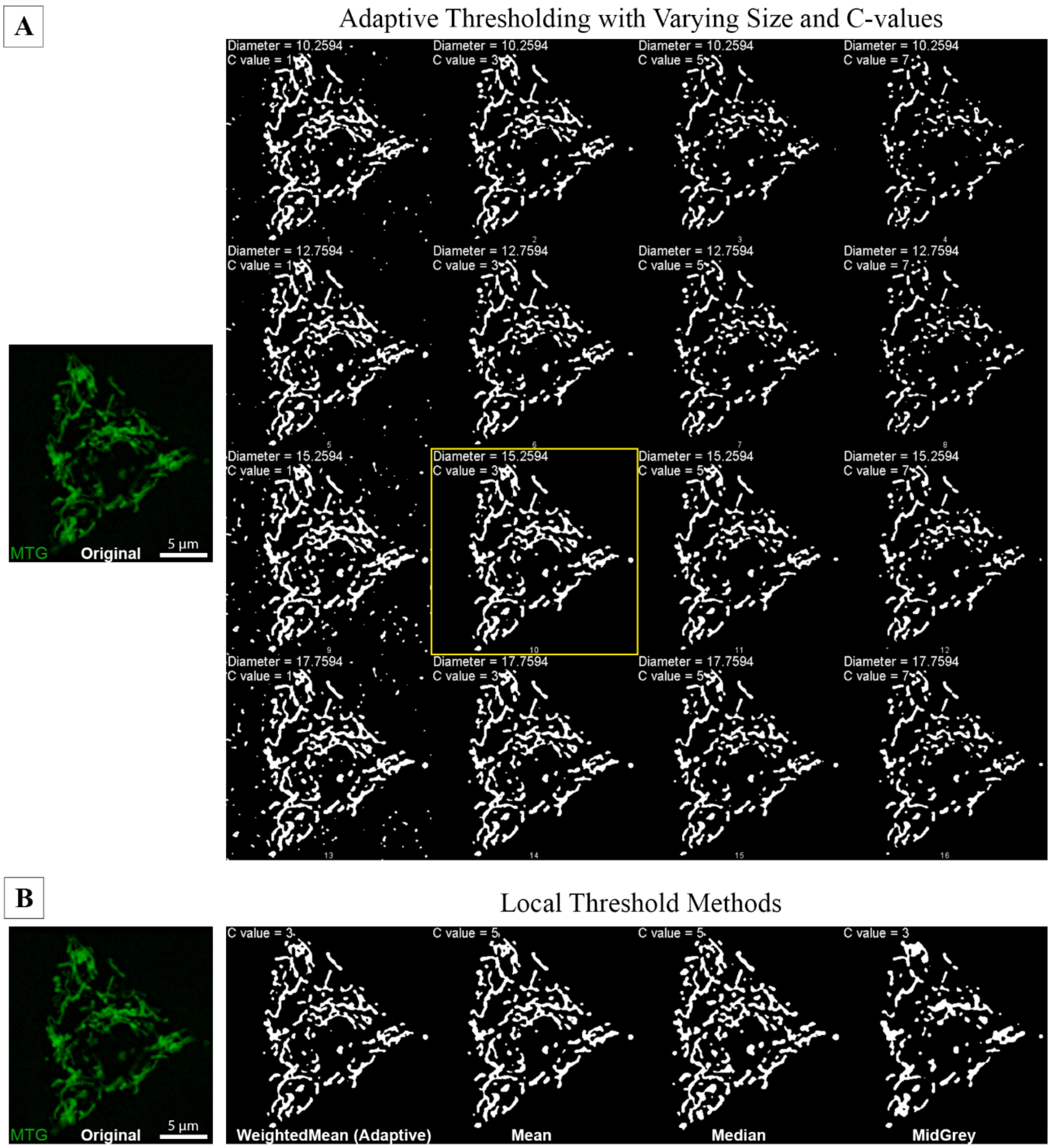


**Figure S2: Comparison and optimization of local thresholding algorithms in ImageJ/Fiji.** Local threshold algorithms identify positive pixels based on analysis of image sub-regions and take two parameter inputs - block size and C-value, both of which require optimization. **(A)** An example of the parameter optimization process, shown here for Adaptive thresholding. *Left* – The original image of an islet cell stained with MitoTracker Green (same image as used in Figure S1). *Right* – We generated a montage of different combinations of block size (diameter in pixels) and C-values to determine the optimal values for faithful segmentation and minimal detection of noise. In this example, we identified a block size of 1.25 µm (15.2594 pixels) and a C-value of 3 as optimal (shown in the yellow box). The block size relates to the diameter of the objects of interest. Mitochondrial diameters vary between 0.3 and 1 µm, suggesting that the optimal block size is slightly larger than the upper end of this mitochondrial size range. **(B)** Comparison of different local threshold algorithms applied with their individually optimized block size and C-values. The empirically determined best block size was consistent across the methods (1.25 µm or 15.2594 pixels), but the optimal C-value varied. The Weighted Mean (Adaptive thresholding) and Mean methods were judged to best preserve structural detail, but in general Adaptive thresholding captured slightly less noise and was therefore chosen for further comparisons and pipeline development.

### Chaudhry et al. – Supplemental Figure 3

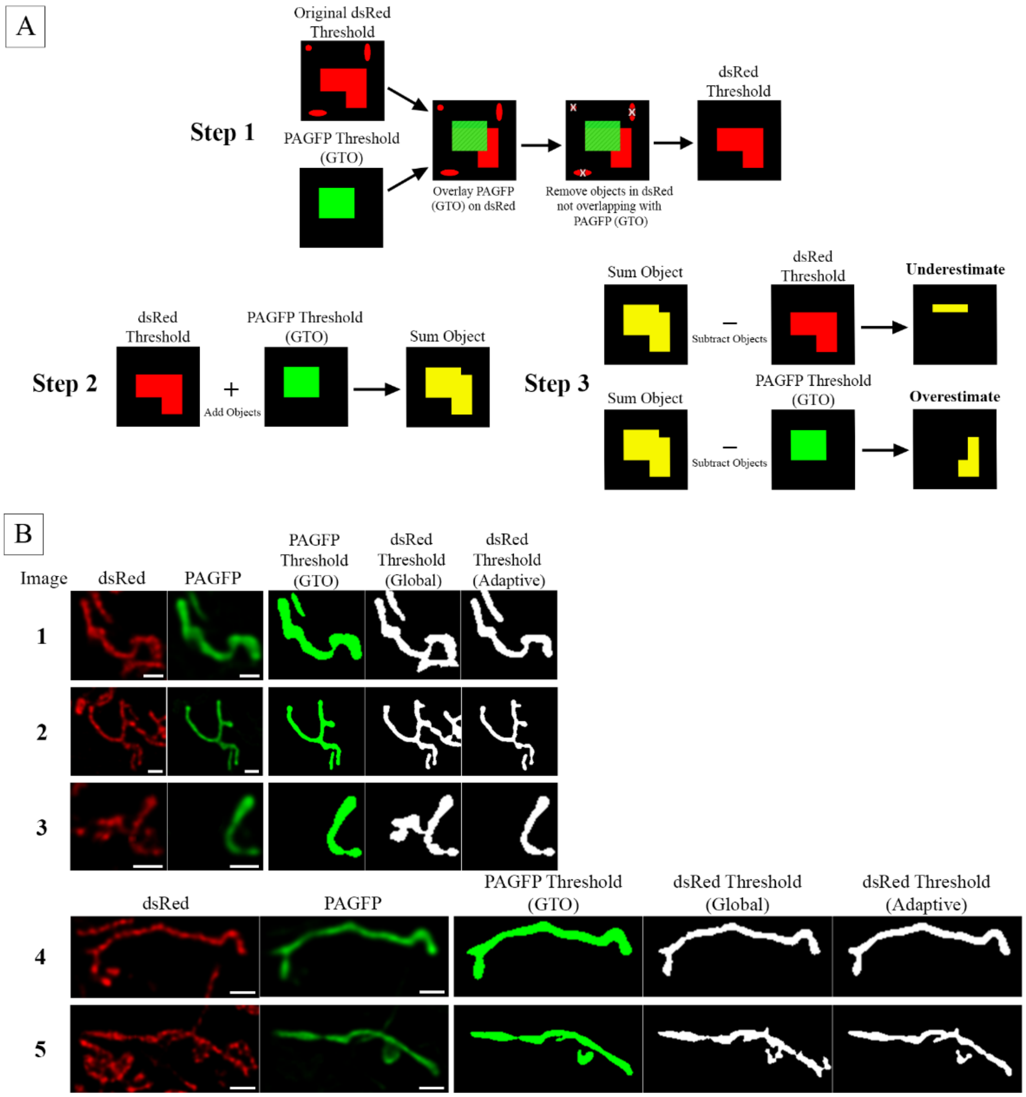


**Figure S3: Quantitative comparison of Global and Adaptive thresholding algorithms in ImageJ/Fiji**. MIN6 cells were co-transfected with Mito-dsRed and Mito-PAGFP to evaluate the accuracy of Global and Adaptive thresholding algorithms. PAGFP-labelled mitochondria served as 'ground truth objects' (GTO) to which the outcome of thresholding was compared. The Global and Adaptive threshold algorithms were applied to the dsRed-labeled mitochondrial population and their performance was quantified by how much the resulting objects overestimated and underestimated the GTO. **(A)** Schematic illustration of how the degree of overestimation and underestimation was determined. *Step 1*: Any objects in the original dsRed threshold (shown in red) that did not connect or overlap with the GTO (shown in green) were identified and removed. *Step 2:* A Sum Object (shown in yellow) was generated by merging the GTO and the remaining dsRed threshold object using the binary "Add" command in the ImageJ/Fiji Image Calculator. *Step 3:* The underestimated area was derived by subtracting the dsRed threshold object from the Sum Object using the binary "Subtract" command. Similarly, the overestimated area was derived by subtracting the GTO from the Sum Object. Finally, the degree of overestimation (OE) and underestimation (UE) was calculated as a percentage of the GTO: OE = OE Area / GTO Area, and UE = UE Area / GTO Area. **(B)** The 5 different test images used to generate the data shown in Figure 2C. In each set of images, the first and second columns show the raw dsRed and PAGFP channels, respectively. The third column shows the GTO, which was identified by Global thresholding of the PAGFP channel. The dsRed threshold objects that overlapped, or connected, with the GTO are shown in white in rows 4 (Global threshold) and 5 (Adaptive threshold). The degree of over- and under-estimation was calculated by comparing these objects to the PAGFP-identified GTO, as detailed in panel A. The over- and under-estimation results are summarized in Figure 2C. Scale bar = 1μm.

### Chaudhry et al. – Supplemental Figure 4

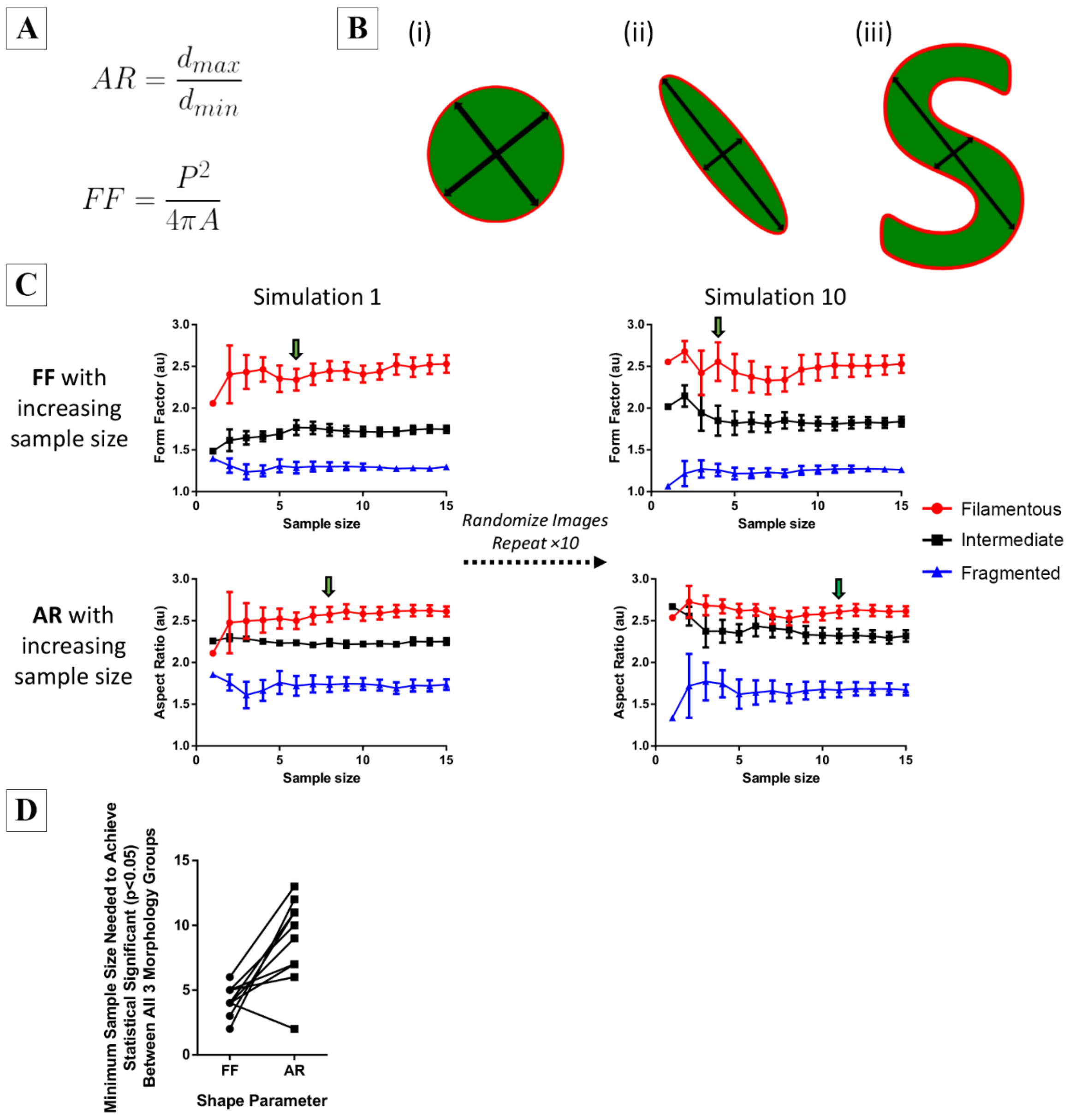


**Figure S4 (Related to Figures 4 and 5): Comparison of Form Factor (FF) and Aspect Ratio (AR) for detection of differences in mitochondrial shape. (A)** Mathematical definitions of FF and AR; $d_{max}$ = maximum length of major axis of fitted ellipse, $d_{min}$ = minimum length of minor axis of fitted ellipse, P = perimeter, A = area. **(B)** Illustration of how FF and AR change between 3 theoretical shapes: i) For a perfect circle both FF and AR are equal to 1.0. ii) As the object elongates both FF and AR increase. iii) Example of a more irregular shape where FF continues to increase but AR is unchanged from the shape in (ii) because the minor and major axes remain the same. **(C)** Simulated experiments comparing the minimum sample size needed for FF and AR to detect significant differences between fragmented, intermediate or filamentous mitochondrial morphologies. In each simulation, mean FF and AR were calculated based on an increasing number of images that were chosen randomly from each morphological category in the test image-set (see Figures 4 & 5). The process was repeated 10 times and simulations 1 and 10 are shown for illustration. Arrows indicate the minimum sample size at which all the morphological groups differed significantly by Repeated One-way ANOVA ($p < 0.05$). **(D)** Summary of the minimum sample sizes needed for FF and AR to distinguish the three morphological categories in each simulation. In 9 out of 10 tests FF required lower sample size than AR, strongly indicating it is a more sensitive mathematical descriptor of mitochondrial shape in 2D analyses.

**Chaudhry et al. – Supplemental Figure 5**

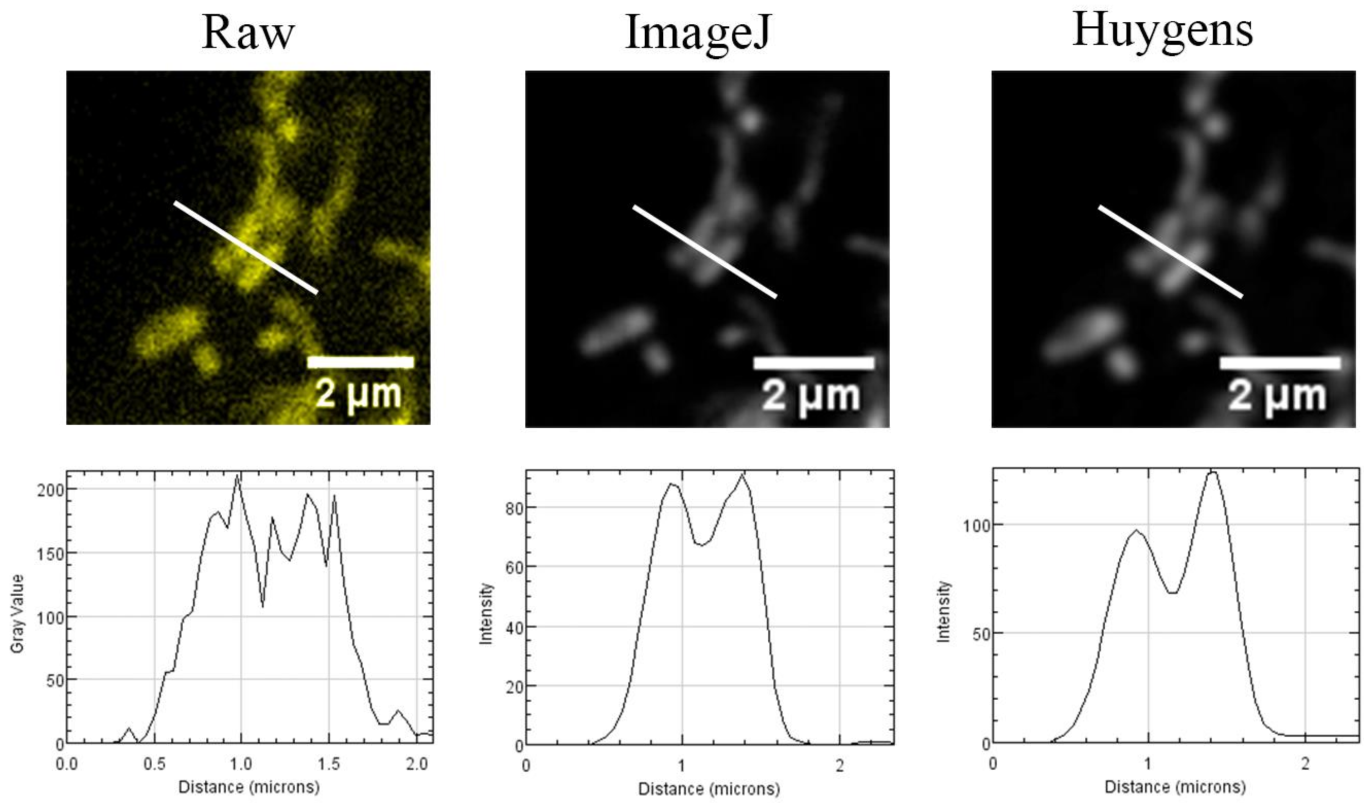


**Figure S5 (Supplement to Figure 7): Demonstration of the improvement in signal quality and contrast between adjacent objects by deconvolution.** *Top row* – A sub-region of the image used in Figure 7 showing two adjacent mitochondrial tubules before deconvolution (Raw) and following deconvolution using the DeconvolutionLab2 module (ImageJ) or Huygens Professional (Huygens). See Figure 9 and Materials & Methods for details. *Bottom row* – Comparison of intensity profiles across the white line indicated in the top images show that deconvolution increases the contrast between adjacent fluorescent objects.

**Chaudhry et al. – Supplemental Figure 6**

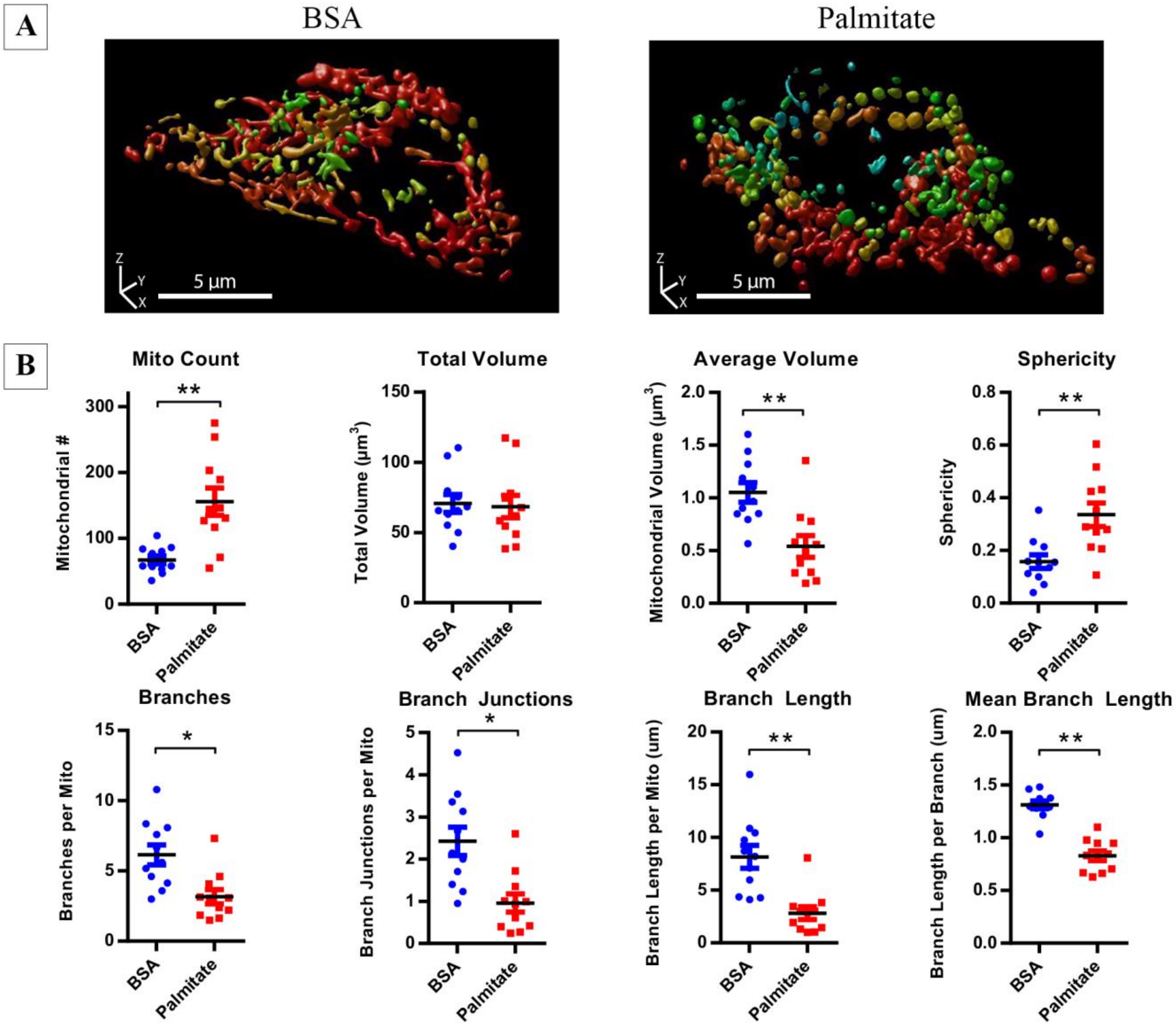


**Figure S6**. **Three-dimensional characterization of lipotoxicity-induced mitochondrial network disruption in MIN6 cells**. Mito-YFP expressing MIN6 cells were treated either with BSA control media or 1.5 mM palmitate for 6 hours before confocal image stacks were acquired of the whole cells. **(A)** 3D renderings (produced in Huygens) of representative cells treated with BSA (left) or palmitate (right). **(B)** Quantitative 3D analysis and comparison of mitochondrial morphology and network connectivity demonstrate a marked fragmentation of the mitochondrial network in palmitate-treated cells. All data are represented by mean ± SEM. *p<0.05, **p<0.01, ***p<0.001 as determined by two-tailed Student's t-test; n=11 cells in each treatment condition.

### Chaudhry et al. – Supplemental Table 1

**Table S1**: Effects of deconvolution using either Huygens or ImageJ software on image restoration and quantitative 3D analysis of the images depicted in Figure 7.

| Property | | Raw | ImageJ | Huygens |
|---|---|---|---|---|
| Image Restoration | | | | |
| | Axial Reduction | - | 74% | 52% |
| | XZ to XY ratio | 2.46 | 1.81 | 1.20 |
| 3D Analysis | | | | |
| | Total Volume | 83.0 | 55.85 | 50.1 |
| | Mitochondria Number | 50 | 76 | 78 |
| | Mean Mitochondrial Volume | 1.66 | 0.73 | 0.64 |
| | Surface Area | 659.4 | 477.9 | 448.9 |
| | Sphericity | 0.12 | 0.15 | 0.21 |
| | Branch Number / Mito | 5.6 | 4.1 | 4.4 |
| | Branch Length / Mito | 5.8 | 4.6 | 4.5 |
| | Branch Junctions / Mito | 2.4 | 1.4 | 1.4 |
